# Differential spatial regulation and activation of integrin nanoclusters inside focal adhesions

**DOI:** 10.1101/2023.12.16.571970

**Authors:** Sarah Keary, Amaris Guevara-Garcia, Nicolas Mateos, Felix Campelo, Maria F. Garcia-Parajo

## Abstract

α_5_β_1_ and α_v_β_3_ integrins are core components of focal adhesions (FAs) involved in cell attachment, migration, and mechanobiology-dependent processes. Recent works indicate that both integrins organize in nanoclusters inside FAs, with sub-populations of active and inactive β_1_ nanoclusters. However, whether both integrins work in concert or their activities are spatially regulated is not fully understood. Using dual-color super-resolution microscopy (STORM, STED and DNA-PAINT) we show that integrins α_5_β_1_ and α_v_β_3_ and their main adaptor proteins exhibit similar functional nanoscale segregation across all adhesion proteins examined, independent of FA maturation state. Notably, both integrins never mix at the nanoscale indicating that their functions might be spatially regulated. We find a nearly 1:1 relationship between active integrin and adaptor nanoclusters suggesting that coordinated integrin activation occurs via the concurrent engagement of adaptor nanoclusters. Interestingly, α_5_β_1_ nanoclusters preferentially localize at the FA periphery near adaptor nanoclusters, establishing regions of multi-nanocluster enrichment, whereas α_v_β_3_ nanoclusters uniformly distribute throughout FAs. Overall, our results show that adhesion proteins are arranged as modular nanoscale units that distinctively organize inside FAs to spatially regulate integrin activation and function.

## Introduction

Adhesion complexes (ACs) in adherent cells, such as focal adhesions (FAs) and fibrillar adhesions (fAs) are places for cell attachment, mechanosensing and mechanotransduction (1–4). These structures are composed by different integrin receptors that bind to their ligands on the extracellular matrix (ECM), and by a myriad of intracellular adaptors, proteins that dynamically connect integrins to the actin cytoskeleton machinery relaying the transmission of mechanical forces in a bidirectional manner (5–7). Studies over the last two decades have provided a wealth of information focused on understanding these complexes, the proteins they contain, their role in cellular processes, such as mechanosensing, migration and proliferation, as well as how influential their regulation is in developmental and pathogenic processes (8–15).

Integrins α_5_β_1_ and α_v_β_3_ are transmembrane heterodimeric proteins critical for the formation of ACs. Both α_5_β_1_ and α_v_β_3_ bind to the RGD peptide motif of certain ECM components, such as fibronectin (16–18), and are linked to the cell’s cytoskeleton via many of the same adaptor proteins (19–22). The apparent redundancy of different integrins binding to the same ligand and same adaptor proteins is intriguing and has been the subject of intense research over the past years. Earlier studies on primary fibroblasts showed that these integrins segregate in different adhesion structures, indicating different roles, with α_5_β_1_ being enriched in fAs (23), adhesive sites at central regions of the cell, and α_v_β_3_ being confined to focal complexes at the base of cellular protrusions, and to FAs, which anchor contractile stress fibers at the periphery of the cell (24). However, more recent super-resolution microscopy studies showed that both types of integrins are simultaneously present inside FAs (25). Within FAs, α_5_β_1_ has been proposed as the integrin responsible for adhesion strength while α_v_β_3_ enables mechanotransduction, as cells exposed to α_5_β_1_ blocking antibody exhibited weaker attachment than those whose α_v_β_3_ integrins were blocked (5). These results are also aligned to *in vitro* reconstitution experiments, that have assessed the catch-bond behavior of both integrins with fibronectin. In this type of bond, the lifetime of the bond increases as the applied force increases, up to a certain threshold, beyond which the bond ruptures. The rupture force for α_5_β_1_ binding to its ligand has been found to be higher (∼10–30 pN) than that of α_v_β_3_ (∼5–25 pN) (26, 27). Nevertheless, single molecule tracking data indicated that α_v_β_3_ integrins are mainly stationary inside FAs and more stably bound to the ligand, favoring adhesion as its main role (25). It has been also shown that α_5_β_1_ is involved in Rho activation during late spreading, enhancing RhoA-mediated actomyosin contractility (24, 28), supporting the idea that α_5_β_1_ is involved in force-mediated adhesion reinforcement. In a *tour-de-force* effort, Fässler and colleagues found that only the simultaneous expression of both integrins enabled cells to sense the rigidity of the ECM and to respond by reinforcing the actomyosin structure (29). These results, supported by others (30), have led to the recent notion that both integrins work in a synergistic manner, exhibiting cooperativity during cell adhesion, migration and mechanosensing, although the mechanisms are far from being fully resolved (30).

The highly dynamic conformational states exhibited by integrins together with their lateral and axial nanoscale organization might play important roles regulating the differential or synergistic functions of different integrins inside FAs. Indeed, it has been shown that integrins can rapidly transition between different conformational states that relate to their activation and binding capacity to their ligands (25, 31, 32). Moreover, in addition to the nanoscale axial segregation of different AC components inside FAs (33), it has been reported that integrins as well as different adaptor proteins also segregate laterally into small nanoclusters inside FAs (34–37). Recent super-resolution studies have further shown that β_1_ integrins segregate in both active and inactive nanoclusters inside FAs (36) and single-molecule tension measurements uncovered distinct sub-populations of load bearing integrins and a high fraction of unengaged integrins (38, 39). Together, these results suggest that at any given time only a fraction of integrins might be active inside FAs and engaged with their adaptor partners. Such a dynamic scenario would allow integrins to differentially interact with their environment and possibly diversify their functions. As α_5_β_1_ and α_v_β_3_ have different activation states and are both contained in FAs, it is plausible that their spatial regulation and activation state inside FAs provides a physical mechanism to tune their functions. However, such studies have not been reported yet.

Here, we combined dual-color super-resolution microscopy together with robust analytical tools to study a number of endogenous FA proteins on human fibroblast cells. Specifically, we imaged one of the two integrins (α_5_β_1_/α_v_β_3_), their full population or active conformations, with each of three key adaptors proteins, paxillin, talin and vinculin. Our results show that each of these proteins form segregated nanoclusters of remarkably similar size and molecular density inside FAs, and interestingly, also on the membrane outside of ACs. Moreover, we find sub-populations of active and inactive β_1_ and β_3_ nanoclusters inside FAs, which spatially segregate from each other and provide evidence for a nearly 1:1 relationship between talin, vinculin and active integrin nanoclusters. Interestingly, we find that α_v_β_3_ nanoclusters organize randomly inside FAs regardless of their activation state, whereas α_5_β_1_ and in particular the active conformation of β_1_ nanoclusters preferentially organize at the edges of FAs. Overall, we propose that the nano- and meso-scale organization of α_5_β_1_ and α_v_β_3_ integrins and their distinct lateral segregation and activation state are key factors in the differentiation of their roles within FAs.

## Results

### Proteins of the ACs organize as universal nanoclusters across regions of the basal membrane

We employed dual-color super-resolution microscopy based on stochastic optical reconstruction microscopy (STORM) to determine how adhesion proteins (integrins and key adaptor proteins) spatially organize at the nanoscale. Human fibroblasts (HFF-1 cells) were seeded on fibronectin (FN)-coated glass slides for 24 h, after which the cells were processed for immunofluorescence. One integrin (either the total population of α_5_β_1_ or α_v_β_3_, or the active population specific to the β_1_ or β_3_ integrin subunit) was labeled together with one adaptor protein (paxillin, C terminus of talin-1, or vinculin) and then imaged using STORM. We note that while the pan antibodies used to target the full population of integrins are specific to α_5_β_1_ or α_v_β_3_, (with the anti-α_5_ antibody binding the α_5_ subunit, which only dimerizes with β_1_, and the anti-α_v_β_3_ targeting this specific heterodimer), the active antibodies target the β subunits. These β subunits can dimerize with other α subunits, but with our experimental conditions, where FN is the most abundant ligand (see Materials & Methods), the primary integrins being probed are the heterodimers, α_5_β_1_ or α_v_β_3_. Given the huge number of adaptor proteins that have been reported to interact with integrins (19, 20), we chose for paxillin, talin and vinculin since they belong to the core-components of the adhesome (21) and crucially locate at distinct axial layers of functional activity, i.e., paxillin at the signaling layer, while talin and vinculin locate at the force-transduction layer (33).

STORM super-resolution images qualitatively revealed an inhomogeneous distribution for all the proteins examined, forming small and laterally segregated nanoclusters inside FAs (Fig. 1A–D). To quantitatively substantiate these observations, we first used a mask to classify all the single molecule localization events according to their location on the membrane, i.e., FAs, fAs or outside ACs (see Materials & Methods, Fig. S1A-F). To identify whether multiple localization events correspond to nanoclusters we used a density-based spatial clustering of applications with noise (DBSCAN) algorithm (40), which has been extensively used for quantifying protein nanoclustering from single molecule localization images (41–44), including β_1_ integrins inside FAs (36, 45). Since STORM data consist of lists of single-molecule localizations and each labeled protein can be localized multiple times due to fluorophore blinking (46) resulting in pseudoclusters (i.e., non-biological artefactual clusters (41)), we first assessed the number of localizations detected due to the multiple blinking events of a single fluorophore. For this, we followed an approach similar to that reported by Spiess et al. to generate distributions of the number of localizations obtained from isolated fluorescent spots on glass next to the cells likely corresponding to individual antibodies (Abs), for both channels (Fig. 1E, Materials & Methods). We note that providing quantification on the FN-coated glass substrate next to the cells guarantees identical experimental and imaging conditions, thus avoiding potential differences in blinking photophysics as compared to calibration measurements performed solely on glass (i.e., without the cells). By following this approach, one can determine the number of localizations corresponding to individual Abs and infer whether the clustered localizations observed correspond to individual or multiple proteins. The median values of the distributions for individual Abs were 5 localizations for Alexa 405-Alexa 647-conjugated Abs (Channel 1) and 3 localizations for Cy3–Alexa 647-conjugated Abs (Channel 2) (Fig. 1E). Moreover, to further minimize the effect of fluorophore blinking on cluster identification, we imposed an additional threshold of 10 localizations as commonly used in the field (36, 45). Thus, any cluster identified by DBSCAN with 10 or more localizations was considered as a cluster containing more than one protein (see Materials & Methods). DBSCAN classified between 70% to 80% of the localizations of each of the proteins as belonging to nanoclusters, regardless of whether they were located inside FAs or on regions of the membrane outside ACs (Fig. S1G-I). Notably, the number of localizations per nanocluster and their size were similar across the different adhesion proteins (integrins and adaptors) inside FAs, with a median of ∼20 localizations per nanocluster, likely corresponding to a few copies of the protein, and a diameter of ∼ 40–45 nm (Fig. 1F, G).

**Figure 1.**
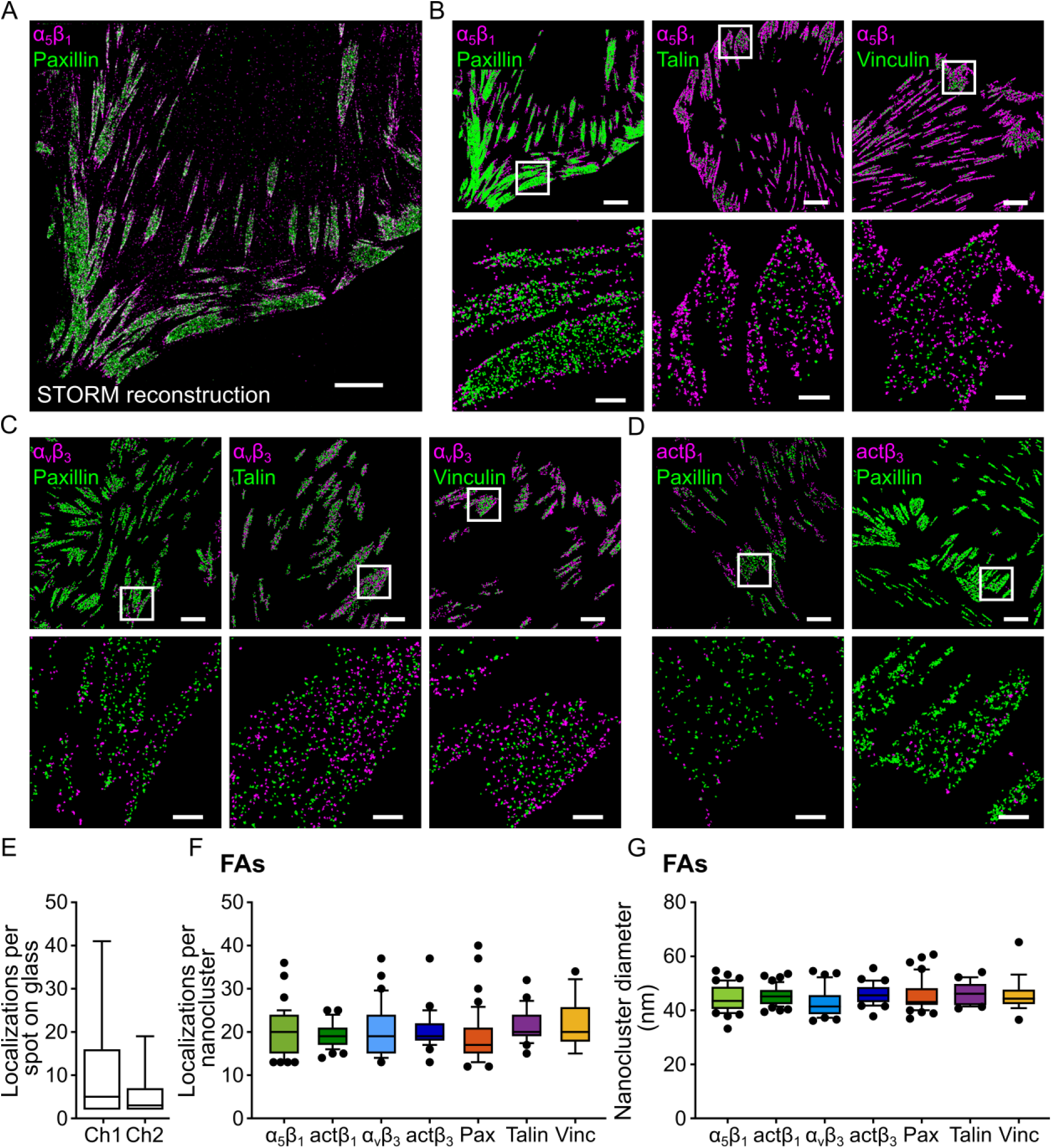
Proteins of FAs organize as universal nanoclusters of similar size and containing a similar number of proteins. (A) Representative dual-color STORM image of an HFF-1 cell fixed after 24 h on FN and labeled for α_5_β_1_ (magenta) and paxillin (green). Scale bar: 5 μm. (B) Representative images of all the adhesions identified for α_5_β_1_ with paxillin (left), talin (middle) and vinculin (right), with zoomed-in regions shown below each panel. Scale bar top row: 5 μm, bottom row: 1 μm. (C) Representative images of all the adhesions identified for α_v_β_3_ with paxillin (left), talin (middle) and vinculin (right), with zoomed-in regions shown below each panel. Scale bar top row: 5 μm, bottom row: 1 μm. (D) Representative images all the adhesions identified for of active β_1_ (act β_1_) with paxillin (left), active β_3_ (act β_3_) with paxillin (right), with zoomed-in regions shown below each panel. Scale bar top row: 5 μm, bottom row: 1 μm. (E) Distribution for the number of localizations per secondary antibody on glass next to cell regions for both of the secondary antibodies used in this study, goat anti-mouse Alexa 405–Alexa 647 (Ch1) and goat anti-rabbit Cy3–Alexa 647 (Ch2). The median number of localizations per spot corresponds to: 5 for Alexa 405-Alexa 647-conjugated antibodies and 3 for Cy3–Alexa 647-conjugated antibodies (see Materials & Methods). (F, G) Box-and-whisker plots showing the number of localizations per nanocluster (F) and the nanocluster diameter (G) for the proteins in this study. The box represents the range from the 25^th^ to the 75^th^ percentiles and whiskers from 10^th^ to 90^th^ percentiles, with the median indicated by a horizontal line. Points outside these whiskers correspond to outliers. Individual points (points within the whiskers not shown) correspond to the median value over all nanoclusters for each cell (for N values, i.e., number of independent experiments, see SI Table S5). No statistical differences were found using the multi comparison one-way ANOVA test, where not significant (ns) corresponds to p>0.05.

Similar results in terms of diameter and number of localizations per nanocluster were obtained when performing the analysis outside ACs (Fig. S2). In addition, the nanocluster characteristics were maintained when specifically labeling the active integrin (β_1_ or β_3_) populations (Fig. 1 D, F, G). These results are in good quantitative agreement with previous reports using STORM showing that both active and inactive integrin β_1_ nanoclusters have a median size of 40 nm and 35 nm, respectively, as well as comparable number of single molecule localizations per nanocluster (36). Notably, our results provide new insights on the integrin α_v_β_3_ and the main adaptor proteins, paxillin, talin and vinculin, by indicating that the organization into laterally segregated nanoclusters of similar size and number of molecules is maintained across different protein types. Moreover, such nanoscale organization appears to be independent of whether these adhesion proteins partition inside FAs or outside ACs (Fig. S2).

Given the remarkable similarity in terms of nanoclustering size and number of localizations amongst the different proteins investigated and regardless of their location (i.e., inside or outside FAs), we performed super-resolution imaging using two additional and different methodologies, stimulated emission depletion (STED) microscopy (47, 48) and DNA-points accumulation for imaging in nanoscale topography (DNA-PAINT) (49). Because STED and DNA-PAINT rely on different physical mechanisms (fluorescence depletion via stimulated emission depletion in STED, and kinetic binding of short dye-labeled oligonucleotides to their complementary target strands in the case of DNA-PAINT) as well as different technical implementations, they constitute excellent controls to validate the results obtained by STORM. Moreover, albeit DNA-PAINT is a single molecule-based localization method similar to STORM, it is not influenced by fluorophore blinking (50), providing an additional control to our STORM results. Both STED and DNA-PAINT images confirmed nanoscale clustering of the proteins investigated, both inside and outside FAs (Fig. S3, Supplementary Text 1, and Fig. S4, Supplementary Text 2), robustly validating our STORM results.

To further exclude any potential bias in quantifying the extent of nanoclustering as obtained by DBSCAN, we implemented an algorithm based on Voronoi-tessellation (41) and re-analyzed all the STORM data (Supplementary Text 3). This analysis showed similar nanoclustering size and number of localizations per nanocluster as compared to DBSCAN (Fig. S5). Altogether, our experimental results obtained using three distinct super-resolution modalities and by analyzing the images using two different clustering algorithms render comparable sizes in terms of nanoclustering for all the proteins investigated, indicating that proteins of ACs organize as universal nanoclusters of similar size and number of proteins, regardless of whether these nanoclusters are located inside or outside FAs. Nevertheless, we note that quantifying the actual stoichiometry of the nanoclusters, i.e., number of proteins per nanocluster based on fluorescence strictly requires a 1:1 labeling between the protein and the fluorophore which is highly challenging, in particular when employing super-resolution approaches (51–53). Therefore, our relative comparison of nanoclustering among the different proteins investigated is exclusively based on the number of single molecule localizations contained in each nanocluster (both for STORM and DNA-PAINT) which is a fair approach since we always use the same reporter fluorophore and maintain similar excitation conditions throughout our experiments.

### α_5_β_1_ and α_v_β_3_ integrin nanoclusters differentially organize inside FAs

Our results so far indicate that the two major integrins involved in FAs form nanoclusters of similar size and molecular content. To enquire whether they act in concert inside FAs by associating at the nanoscale, we performed dual color STORM imaging of the total populations of α_5_β_1_ and α_v_β_3_ integrins. For this, we used a rabbit anti-α_5_ antibody (Table S2) to enable simultaneous labeling with the mouse anti-α_v_β_3_ antibody that we used throughout this study. Visual inspection of the images already suggests that α_5_β_1_ and α_v_β_3_ do not physically overlap at the nanoscale (Fig. 2A). To robustly quantify these observations, we calculated the edge-to-edge distance between the nearest-neighbor (EED-NN) integrin nanoclusters (i.e., closest α_5_β_1_ to α_v_β_3_ and closest α_v_β_3_ to α_5_β_1_) (see Materials & Methods). Here, EED-NN values below zero correspond to physically overlapping α_5_β_1_ and α_v_β_3_ nanoclusters, EED-NN values above zero to spatially segregated nanoclusters, and EED-NN = 0 corresponds to contiguous α_5_β_1_ and α_v_β_3_ nanoclusters, i.e., next to each other. Quantification of the images confirmed that α_5_β_1_ and α_v_β_3_ do not intermix at the nanoscale as the percentage of EED-NN values below zero is negligible and comparable to values obtained from simulations of random organization of both integrins (Fig. 2B). Thus, α_5_β_1_ and α_v_β_3_ integrins organize as distinct nanoclusters that spatially segregate from each other inside FAs.

**Figure 2.**
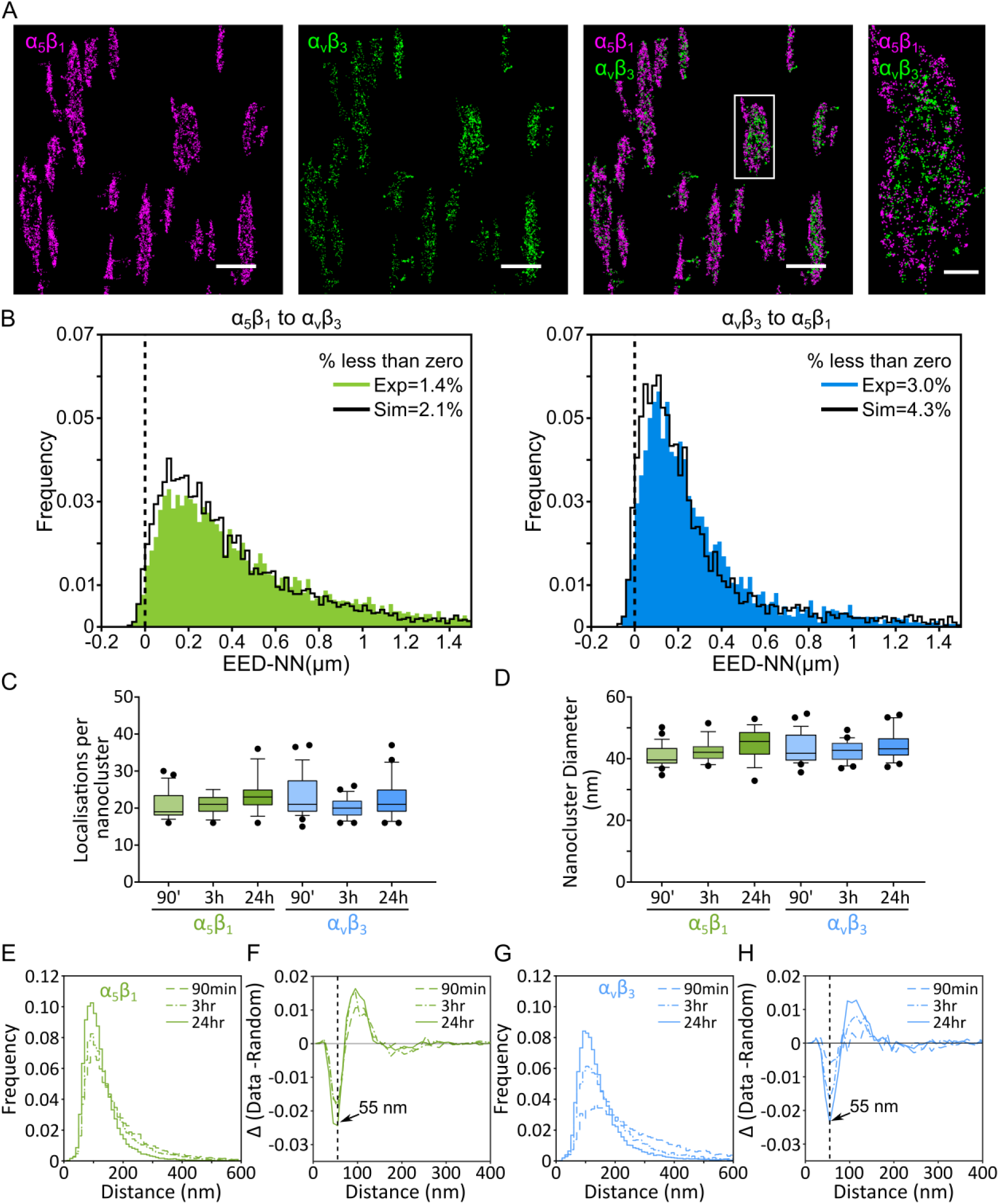
α_5_β_1_ and α_v_β_3_ integrin nanoclusters differentially organize inside FAs. (A) Representative dual-color STORM image of the total populations of α_5_β_1_ (magenta) and α_v_β_3_ (green) integrins inside FAs. The panel on the far-right side corresponds to a zoomed-in region of a single FA (white box) showing spatially segregated α_5_β_1_ and α_v_β_3_ nanoclusters. Scale bars: 2 μm, and 500 nm for the zoomed-in region. (B) Histogram distribution for edge-to-edge distance between nearest neighbors nanoclusters (EED-NN), from α_5_β_1_ to α_v_β_3_ (A, green) and α_v_β_3_ to α_5_β_1_ (B, blue). (C-D) Nanocluster properties inside FAs for the integrins α_5_β_1_ (green), and α_v_β_3_ (blue) as a function of seeding time. (C) show the number of localizations per nanocluster;(D) shows the nanocluster diameter. Only the pairs with statistical differences are marked. *, p<0.05; **, p<0.01; ***, p<0.001; one-way ANOVA. (E-H) Histograms of the experimental NND distributions for the α_5_β_1_ (green) (E) and α_v_β_3_ (blue) (G) integrins in FAs, for the designated cell spreading times. Histograms for the random simulations have been omitted for the sake of visual clarity. (F,H) ΔNNDH-plots for the subtraction histograms for simulated data from experimental for the α_5_β_1_ (F) and α_v_β_3_ (H) integrins in FAs at the three different seeding times (90 min, 3 h and 24 h).

Since FA formation is a dynamic process where mechanical forces and FA maturation increase over several hours, we thought to investigate whether the nanoscale organization and distribution of both integrins inside FAs are sensitive to FA maturation. For this, we performed STORM imaging as described above on cells seeded for 90 min and 3 hours and quantified the number of localizations per nanocluster (Fig. 2C) and nanocluster diameter (Fig. 2D) for both integrins and, compared the results to 24 hours cell seeding. No differences in nanocluster properties were observed for any of the two integrins suggesting that integrin nanoclustering is independent of spreading times and FA maturation. To then investigate how integrin nanoclusters are distributed inside FAs, we performed nearest-neighbor distribution (NND) analysis for both integrins at different cell seeding times (Fig. 2E, G). Since the NNDs are influenced by the nanocluster densities, we compared the experimentally obtained NNDs with those obtained from simulations of randomly distributed nanoclusters considering the experimentally obtained nanocluster density. We then calculated the difference between experimental and simulated data for each NND value, i.e., ΔNND for both integrins at different cell spreading times (Fig. 2F, H). Interestingly, our analysis revealed a clear deviation from random organization inside FAs, with both integrin nanoclusters being segregated from each other at around 55 nm and enriched at distances ∼ 100-150 nm. Strikingly, this physical segregation and enrichment depended on cell spreading time and it was different for both integrins: α_5_β_1_ segregation and enrichment was already established at earlier spreading times (90 min) (Fig. 2F), while α_v_β_3_ nanoclusters organized rather randomly inside FAs at 90 min and progressively established their characteristic spatial segregation at ∼ 55 nm and enrichment at ∼ 120 nm at 24 hours of cell spreading (Fig. 2H). Overall, these results indicate that while integrin nanocluster properties are preserved and maintained regardless of cell spreading times and FA maturation, their distribution and organization inside FAs depends on FA maturation, with α_v_β_3_ lateral ordering being progressively reached at later stages of cell adhesion.

### Lateral organization of adaptors and integrin nanoclusters inside FAs

We next sought to investigate how the different adaptor proteins laterally organize inside FAs beyond their nanoclustering. For this, we first quantified the density of nanoclusters per unit area of the membrane, within FAs (Fig. 3A) and outside ACs (Fig. 3B). As expected, the density of nanoclusters was larger in FAs as compared to outside adhesions (∼5–10-fold increase) (Fig. 3A, B). Adaptor proteins, such as talin or vinculin, are shared between α_5_β_1_ and α_v_β_3_ integrins (21), therefore we compared the density of the full population of integrin nanoclusters (∼36 µm^-2^ in FAs) to the density of adaptor protein nanoclusters (∼10–12 µm^-2^ for talin and vinculin, ∼20 µm^-2^ for paxillin in FAs). These data reveal a significantly higher (∼2–4 fold) density of integrin nanoclusters in comparison to their adaptors and, consequently, a nanocluster mismatch between the number of integrins and adaptor nanoclusters, regardless of whether they partition inside or outside FAs (Fig 3A, B). Since the active integrin conformation is the one engaged with the adaptors (54, 55), we also quantified the density of active integrin nanoclusters (active β_1_ and active β_3_) per unit area inside FAs. Our results showed that only ∼ 30% of the total α_5_β_1_ nanocluster population was in an active conformation, and this percentage was even lower for β_3_ nanoclusters (∼ 20%) (Fig. 3A), indicating that at any given time there is a large percentage of inactive integrin nanoclusters inside FAs. We note that these percentages might be even lower considering that our Abs for mapping the active conformations are not exclusive for α_5_β_1_ and α_v_β_3_, but rather to the total active β_1_ and β_3_ sub-populations.

**Figure 3.**
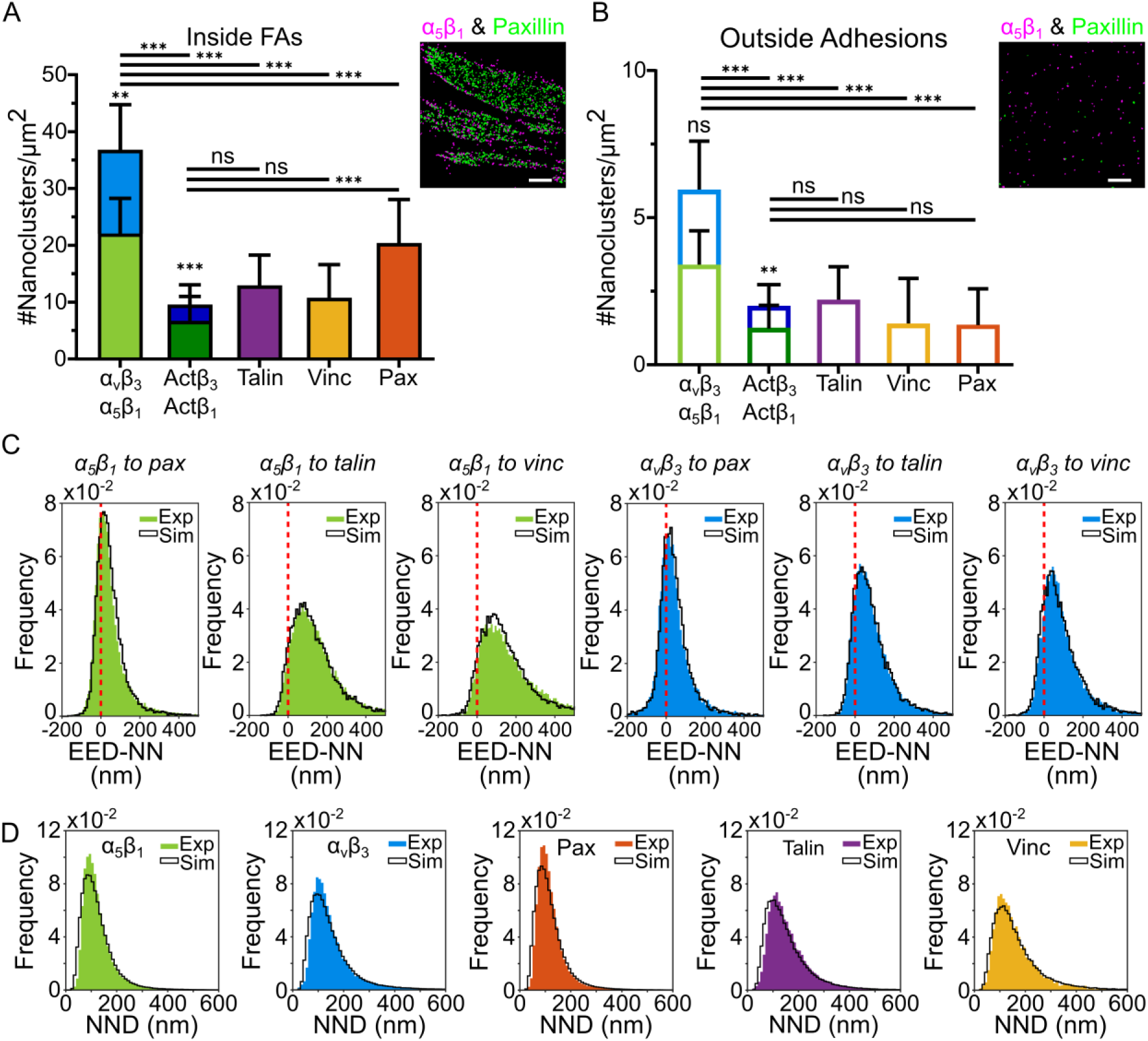
Lateral organization of integrin nanoclusters and adaptor proteins inside FAs. (A, B) Mean number of nanoclusters per unit area inside FAs (A) and on the cell membrane outside adhesions (B), with α_5_β_1_ and actβ_1_ in light and dark green, respectively; and α_v_β_3_ and actβ_3_ in light and dark blue, respectively. The number of integrins nanoclusters per unit area are shown as stacked graphs to allow comparison between the total number of integrin nanoclusters (α_5_β_1_ and α_v_β_3_) and the adaptor proteins. The bars show the mean ± st. dev of the distribution. Statistical significance between data of all proteins was tested using a multi comparison one-way ANOVA, assuming Gaussian distribution of points. The comparison between α_5_β_1_ and α_v_β_3_, (actβ_1_ and actβ_3_) was done with an unpaired t-test, where ns = not significant, p>0.05; *, p<0.05; **, p<0.01; ***, p<0.001. The insets show representative images of nanoclusters for α_5_β_1_ (magenta) and paxillin (green) in FAs (A) and outside adhesions (B). Scale bar: 1 μm. (C) Histogram of the EED-NN values between α_5_β_1_ and the different adaptors (green) and, between α_v_β_3_ and the adaptors (blue). The black line in each plot corresponds to the histogram for the simulated data sets. The dashed red line at zero corresponds to nanoclusters contiguous to each other. Percentage of overlapping nanoclusters: α_5_β_1_-paxillin: exp = 28.4%, sim = 24.4%; α_5_β_1_-talin: exp = 6.7%, sim = 8.1%; α_5_β_1_-vinculin: exp = 6.0%, sim = 7.7%; α_v_β_3_ -paxillin: exp =31.7%, sim=29.2%; α_v_β_3_ -talin: exp =15.7%, sim=16.5%; α_v_β_3_-vinculin: exp =14.9%, sim=17.4%. (D) Histogram of the NND values between nanoclusters of the same protein. The black line in each plot corresponds to the histogram for the simulated data sets. Bin width is 10 nm.

Notably, the total density of active β_1_ and β_3_ integrin nanoclusters matches that of talin and vinculin nanoclusters, and it is only slightly outnumbered by paxillin nanoclusters inside FAs (Fig. 3A), suggesting that these active integrins might be fully engaged to their adaptors. The modest percentage of active β_1_ integrin nanoclusters observed in our data (∼ 30%, Fig. 3A) is in close agreement with a recent report from Spiess *et al*, obtained using a different super-resolution technique (36) and consistent with the high percentage of low-force bearing integrins measured by using single-molecule force sensors (39). In addition to the α_5_β_1_ results, our data also reveal the existence of a sub-population of active β_3_ nanoclusters inside FAs (Fig. 3A). Together, these results strengthen the hypothesis proposed by Spiess et al. of a coordinated mechanism that laterally segregates active and inactive nanoclusters inside FAs (36). Moreover, the differences in expression levels, activation states, and lateral nano- and lateral scale segregation of α_5_β_1_ and α_v_β_3_ nanoclusters inside FAs suggest the existence of a mechanism that defines the distinct functional roles of these two integrins inside FAs.

To get a better understanding on the lateral organization of both types of integrins in relation to their main adaptor proteins, we calculated the EED-NN between integrin nanoclusters and each of the adaptors. Although visual inspection of the dual color STORM images already suggested that integrin nanoclusters are also physically segregated from their adaptor partners (Fig. 1B, C) we used the EED-NN analysis to estimate the degree of colocalization (i.e., spatial overlap) between different nanoclusters. Our results confirm that the large majority of α_5_β_1_ and α_v_β_3_ nanoclusters are laterally segregated from their adaptor partners (Fig. 3C, Fig. S6). Their spatial segregation is essentially determined by the density of nanoclusters within the FAs, as similar EED-NN distributions were obtained from simulations of particles uniformly distributed inside the experimentally obtained FAs, using identical particle densities as to the ones obtained experimentally for the integrin nanoclusters and the different adaptors (Materials & Methods) (Fig. 3C, black lines). Considering the canonical view that integrins inside FAs are bound to paxillin and engaged to talin and vinculin (55, 56), these results might be surprising at first sight, as one would expect a much higher fraction of EED-NN values below zero. Nevertheless, our data showed only a small percentage of active integrin nanoclusters as compared to their total populations (Fig. 3A) so that spatial overlap with talin and/or vinculin would be at best 30% for α_5_β_1_ and 20% for α_v_β_3_ nanoclusters, which under our experimental conditions might be undistinguishable from a random overlap. Notably, using different super-resolution microscopy techniques, others have also found negligible spatial colocalization of integrins with these adaptor proteins (36, 37). Moreover, Dunn and co-workers also found that most ligand-bound integrins inside FAs exist in a minimally tensioned state (< 2 pN) and not dependent on the actin cytoskeleton (39). The results from the Dunn lab are fully in line with the lack of colocalization of integrins to talin and vinculin as we determined here, since these two adaptors mediate the linkage of integrins to the actin cytoskeleton (55, 57). Nevertheless, we would like to mention that we labeled talin at its actin-binding C-terminal region, which in a 2D projection can appear to be at a distance of 100 nm or more from the N-terminal integrin-binding site (58), so lack of spatial overlap does not necessarily indicate lack of direct engagement.

It has been recently proposed that adhesion proteins including integrins organize in a hierarchical fashion inside FAs, forming nanoclusters similar to the ones reported here, that further enrich into compartmentalized regions around 300 nm in size (37). To enquire whether such hierarchical organization is present in our data, we performed NND analysis between nanoclusters of the same proteins (Fig. 3D). In addition, we implemented simulations of uniformly distributed particles inside the experimental FA regions, using the same experimental nanoclusters densities, and made the corresponding NND analysis (Fig. 3D). Our results show that the global organization of all the adhesion protein nanoclusters investigated is highly uniform inside FAs, as the experimental NND distributions were indistinguishable from those of randomly distributed particles (Fig. 3D). In contrast to the recent findings of the Kusumi lab (37), we did not observe any evidence of spatial enrichment of protein nanoclusters inside FAs. Instead, the observed characteristic separation between nanoclusters appears to be solely determined by their density inside FAs.

### Integrin α_5_β_1_ but not α_v_β_3_ nanoclusters are enriched on FA edges

Our previous NND analysis rendered a uniform inter-cluster organization of all adhesion protein nanoclusters inside FAs. Yet, visual inspection of the dual-color STORM images suggests that a significant fraction of α_5_β_1_ nanoclusters preferentially localize on FA edges (Fig. 1B). To quantify these observations, we calculated the shortest-path distance from each α_5_β_1_ nanocluster to the edge of its corresponding FA, i.e., its distance-to-FA-edge, and generated a histogram of these values (Fig. 4A). In addition, we generated a similar histogram from simulations of uniformly distributed α_5_β_1_ nanoclusters on FAs (see Materials & Methods). In general, if the nanoclusters were uniformly distributed throughout the FA, one would expect that the probability of finding a nanocluster at a specific distance to the FA edge decreases with increasing distance. This occurs because there is a larger area along the perimeter of any given shape and thus a higher number of nanoclusters (e.g., for a circular shape, the decrease is linear with the distance to the FA edge). We found that our experimental and simulated data indeed exhibit this general trend, although, notably, a higher percentage of α_5_β_1_ nanoclusters were detected at distances below 100 nm from the FA edges, as compared to a uniform distribution (Fig. 4A and *inset*), indicative of their preference to locate closer to the FA edges. Similar trend was also obtained for talin, and to a lesser extent for paxillin, whereas the distributions of α_v_β_3_ and vinculin were indistinguishable from random (Fig. S7A).

**Figure 4.**
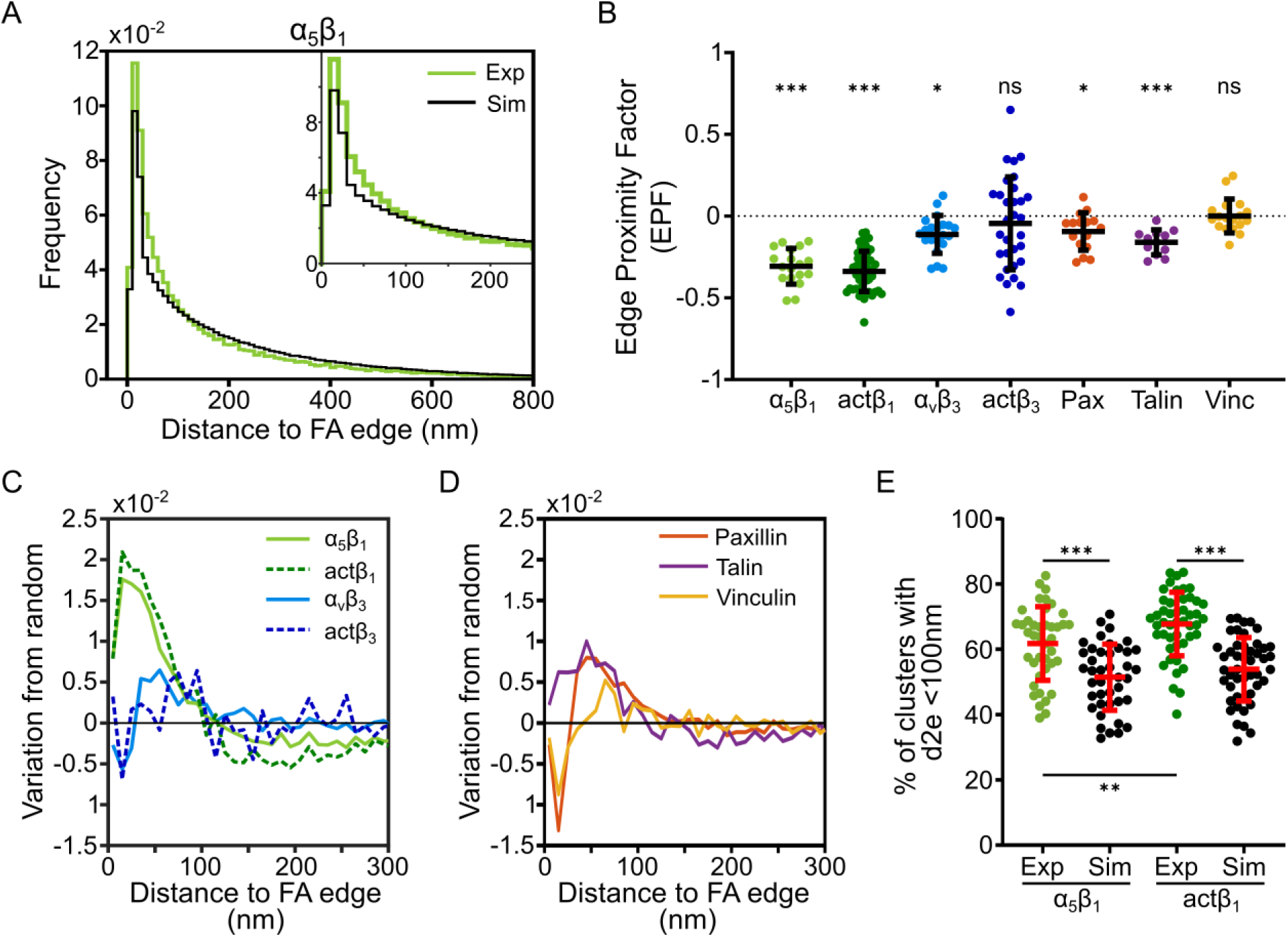
Integrin α_5_β_1_ nanoclusters are preferentially distributed on the edges of FAs. (A) Distribution of distances from the center of mass (CoM) position of each individual α_5_β_1_ nanocluster to its closest FA edge obtained from DBSCAN-analyzed STORM images. The histogram represents the frequency distribution over all cells. Green lines correspond to the experimental data (Exp) and the simulated uniform distribution (Sim) is shown as a black histogram. Bin width is 10 nm. The inset shows the distribution for the first 250 nm. (B) Edge Proximity Factor (EPF) per cell for each of the proteins investigated. Data are shown as median values per individual cell (color dots). The mean ± st. dev. over all the cells are superimposed as black lines. Statistical significance between experimental and simulated data was tested using a paired t-test, where ns = not significant, p>0.05; *, p<0.05; **, p<0.01; ***, p<0.001. (C, D) Variation of the nanocluster distances from the FA edges with respect to random organization, as function of the distance to the FA edges (d2e) for α_5_β_1_ and α_v_β_3_ integrin nanoclusters (C) and for the adaptor proteins (D). (E) Percentage of nanoclusters proximal to FA edges (d2e<100nm) for the total and active β_1_ nanocluster population (color dots) together with results from simulations of random organization (black dots). Each dot corresponds to the median value per cell, each cell containing multiple FAs. The mean ± st. dev. over all the cells is superimposed as red lines. Statistical significance between experimental and simulated data was tested using a paired t-test, and to test the significance between α_5_β_1_ and actβ_1_ the unpaired t-test was used, assuming a Gaussian distribution of data, where ns = not significant, p>0.05; *, p<0.05; **, p<0.01; ***, p<0.001.

To better assess the extent of nanocluster enrichment at FA edges at the single cell level, we computed the median values of the experimentally obtained distance-to-FA-edge distributions per cell (over multiple FAs) and those corresponding to the simulated data. We then defined the edge-proximity-factor (EPF) as the relative difference between experimental and simulated distributions, where negative EPF values correspond to preferential nanocluster enrichment at FA edges, positive EPF values to nanocluster depletion at FA edges, and EPF = 0 corresponds to random location of nanoclusters with respect to the FA edges (Fig. 4B). Moreover, to statistically determine whether the distribution of EPF values was different from zero (the EPF value expected from a random distance-to-the-FA-edge distribution) we ran a paired Student’s t-test for each protein comparing the experimental with the corresponding simulated distributions. This analysis showed that most proteins exhibit some degree of enrichment at the edges of the FAs (i.e., negative EPF values), although their magnitude was markedly different amongst them (Fig. 4B). The highest enrichments at the FA edges were found for the full population of α_5_β_1_ nanoclusters and the active β_1_ sub-population (∼30% closer to the FA edge than if uniformly distributed) and for talin, whereas the enrichment at the FA edges of the total α_v_β_3_ population and paxillin was much more modest. Notably, the active β_3_ sub-population and vinculin showed no preferential enrichment at the FA edges (Fig. 4B).

To establish the characteristic distances at which nanocluster enrichment occurs with respect to the FA edges, we next computed the difference between the experimental and simulated distributions, i.e., the variation from random, as a function of the distance to the FA edges, for each of the examined proteins (Fig. 4C, D). Here, positive values correspond to local nanocluster enrichment at a given distance to the FA edge, whereas negative values indicate nanocluster depletion. Our results show a clear difference between both integrins, with α_5_β_1_ nanoclusters being enriched close to the edge of FAs (Fig. 4C, continuous green line) whereas α_v_β_3_ nanoclusters showed a rather uniform distribution, regardless of their distance to the FA edges (Fig. 4C, continuous blue line). Moreover, the preferential enrichment of β_1_ but not of β_3_ nanoclusters at the FA edges was retained when specifically analyzing the active integrin conformations (Fig. 4C, dashed green and blue lines, respectively).

We next investigated how adaptor protein nanoclusters distribute with respect to the FA edge using a similar approach. We found that both talin and paxillin nanoclusters showed enrichment in a region between 50–150 nm from the FA edges, although somewhat less pronounced as compared to α_5_β_1_ nanoclusters (Fig. 4D), whereas vinculin nanoclusters showed no preference to reside proximal to the FA edges. Finally, we extracted the percentage of the total α_5_β_1_ and active β_1_ nanoclusters that reside within distances shorter than 100 nm from the FA edges and compared the results to those from simulations of random organization (Fig. 4E). Interestingly, a significantly larger percentage of active β_1_ nanoclusters as compared to the total α_5_β_1_ nanocluster population and to the simulations, locate at the periphery of FAs (Fig. 4E).

Together, these results strongly indicate that integrin nanoclusters spatially organize in a rather distinct fashion within FAs, with α_5_β_1_ nanoclusters localizing preferentially at the edges of the FAs and α_v_β_3_ nanoclusters distributing randomly inside FAs. Moreover, the preferential enrichment of active β_1_ nanoclusters together with talin at the FA edges suggests that β_1_ activation is spatially regulated inside FAs.

### Active β_1_ nanoclusters form multiprotein hubs at FA edges

Since activation of integrins involves engagement with their adaptors paxillin, talin and vinculin, we next asked whether the active β_1_ nanoclusters preferentially located at the FAs periphery also reside in proximity to their adaptor proteins. For this we first obtained the inter-nanocluster NND (*i*NND) distributions between active β_1_ nanoclusters and each of the adaptor nanoclusters (Fig. S7B). We calculated the median of each *i*NND distribution and used this value as a threshold to classify pairs of active β_1_-adaptor nanoclusters as being close (i.e., *i*NND < median) or far (*i*NND > median) from each other. Next, we separately computed the distance-to-the-FA edge for the active β_1_ nanoclusters that were found close or far from their partner and compared the results to simulated data (Fig. 5A and *inset*). Having these plots, we then calculated the variation from random (i.e., difference between experimental data and simulations of random organization) as a function of the distance to the FA edge, for both active β_1_ nanoclusters close to their partner and active β_1_ nanoclusters far from their partner (Fig. 5B and Fig. S7C). Surprisingly, this analysis shows that active β_1_ nanoclusters preferentially locate the FA edges regardless of their proximity to talin nanoclusters (Fig. 5B) suggesting that other adaptors aside from talin might contribute to the activation of β_1_ by its engagement with the actin cytoskeleton. Indeed, dual color STED imaging showed co-enrichment of α_5_β_1_ nanoclusters (and the active β_1_ sub-population) together with tensin-3 at the FA periphery (Fig. S8). In contrast, the same analysis on active β_3_ nanoclusters shows a random distribution inside of FAs irrespective of their proximity to talin or other adaptor nanoclusters (Fig. 5C).

**Figure 5.**
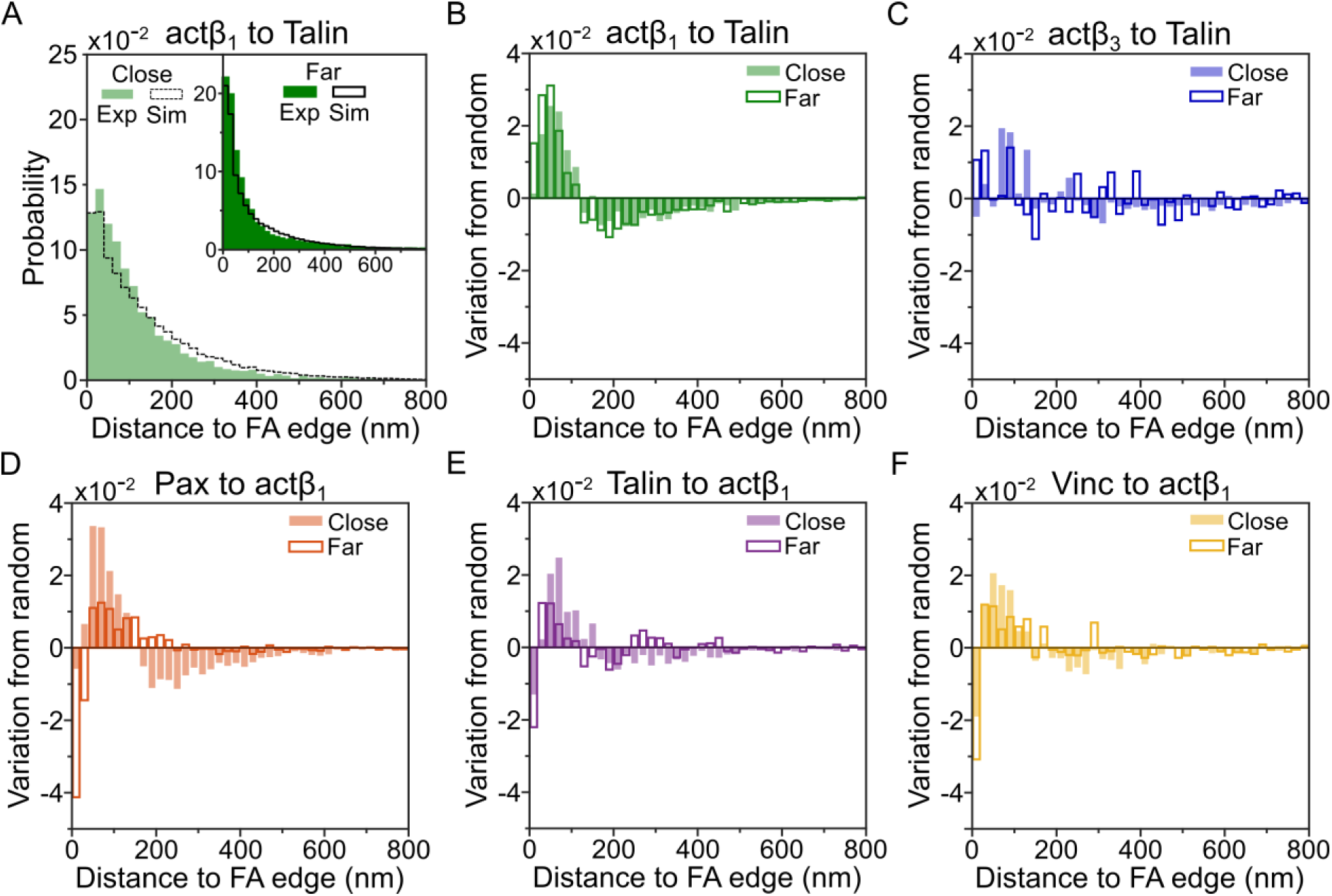
Active β_1_ nanoclusters form multiprotein hubs at the FA edges. (A) Population of active β_1_-talin nanocluster pairs as a function of their distance-to-the-FA edge. The data plotted in light green correspond to active β_1_ nanoclusters located close to talin (i.e., iNND < median), while the inset dark green corresponds to active β_1_ nanoclusters located far from talin (i.e., iNND > median). iNND distributions are shown in Fig. S7B. Black lines are for simulated data sets. Bin width is 20 nm. (B) Variation from random for active β_1_-talin nanocluster pairs as function of their distance-to-the-FA-edge. Filled bars correspond to active β_1_ nanoclusters located close to talin (iNND < median) and empty bars correspond to active β_1_ nanoclusters located far from talin (iNND > median) (C) Similar to (B) but for active β_3_ nanoclusters. (D-F) Variation from random for paxillin (D), talin (E), vinculin (F) active β_1_ nanocluster pairs, as function of their distance-to-the-FA-edge, with data segregated in two different distributions according to the pairs proximity. Bin width is 20 nm.

Finally, we enquired whether adaptor protein nanoclusters (paxillin, talin, and vinculin) close to active β_1_ nanoclusters also locate closer to the FA edges. Remarkably, we found that all of these adaptor nanoclusters close to active β_1_ nanoclusters are also enriched on the FA edge region (∼150–200 nm from the edge), whereas adaptor nanoclusters far from active β_1_ appear to be more uniformly distributed (Fig. 5D–F). Taken together, our data reveal a differential mesoscale organization between active β_1_ and β_3_ nanoclusters inside FAs, with active β_1_ nanoclusters being enriched at the periphery of FAs alongside adhesion adaptor nanoclusters, while active β_3_ nanoclusters remain uniformly distributed throughout FAs, lacking peripheral enrichment.

## Discussion

Here, we have combined dual-color super-resolution optical microscopy together with a robust set of analytical tools to provide insights on the lateral spatial organization of different adhesion proteins inside FAs. Using three different super-resolution imaging strategies, STORM, DNA-PAINT and STED we show that both α_5_β_1_ and α_v_β_3_ integrins as well as their main adaptors, paxillin, talin and vinculin, form nanoclusters of similar size and molecular content inside FAs. We further validated our clustering analysis using Voronoi-tessellation in addition to DBSCAN rendering comparable results in terms of nanoclustering size and content. Moreover, this organization was independent on the membrane location, i.e., inside or outside FAs, or FA maturation, as cells inspected at different spreading times exhibited comparable nanoscale organization of these adhesion proteins.

Earlier reports showed that nascent adhesions (NAs) are formed due to integrin nanoclusters that either rapidly disassemble at the plasma membrane without force or further mature into FAs in response to mechanical forces (35, 59). Our results now show that even in the presence of mechanical forces (i.e., inside FAs), integrin nanoclusters remain segregated from each other while maintaining their sizes and number of molecules. Similar integrin nanocluster properties were also found outside adhesions, reinforcing the notion that integrin nanoclustering is force-independent (35). Interestingly, we found that the main cytoplasmic adaptors, paxillin, talin and vinculin, also form nanoclusters of similar size and number of single molecule localizations as the integrins, regardless of whether they locate inside or outside FAs. The presence of adhesion protein nanoclusters outside of FAs is intriguing. However, immobilized (active) integrins have been previously observed outside of FAs (25, 60). We suggest that these isolated nanoclusters may represent transient nascent adhesions forming on the cell membrane. Although super-resolution images do not allow us to directly extract the absolute number of proteins contained in nanoclusters given the inability to guarantee full labelling saturation and primary-to-secondary antibody stoichiometry, amongst others, the similarity in terms of number of single molecule localizations amongst integrins and their adaptor partners obtained from STORM and DNA-PAINT data suggest that these nanoclusters might be stoichiometrically similar. It will be interesting in the future to devise and/or to adapt emerging super-resolution strategies to determine the precise number of molecules contained in each nanocluster (61–63), although it might take a while until these approaches become routinely available to the community. Together, our results strongly indicate that adhesion proteins organize as universal nano-hubs, functioning as modular adhesion units to serve multiple integrin-dependent functions related to adhesion and migration but also to mechanosensing and mechanotransduction.

Consistent with previous results (36), our data revealed that active and inactive α_5_β_1_ nanoclusters coexist inside FAs but are fully segregated from each other. Remarkably, α_v_β_3_ also exhibits similar nanoscale segregation, with total population-to-active integrin ratios indicating a similar segregation between active and inactive β_3_. Furthermore, earlier data from our group demonstrated that the integrin α_L_β_2_ expressed on leukocytes also segregates into active and inactive nanoclusters (64, 65). Thus, lateral segregation between active and inactive integrin nanoclusters may represent a common feature to spatially coordinate integrin activation. How could such cooperativity in integrin nanocluster activation arise? Our data show a nearly 1:1 matching in the number of active integrin nanoclusters, and the main cytoplasmic adaptors talin and vinculin, known to activate integrins by their engagement to the actin cytoskeleton. The availability of a similar number of adaptor nanoclusters containing comparable number of molecules as the integrin nanoclusters suggests that inactive integrin nanoclusters are poised for their collective nanocluster activation, which is achieved upon the concurrent engagement of talin and vinculin nanoclusters (and most probably also paxillin). This mechanism has been shown to operate in NAs in which concurrent recruitment of talin and vinculin prior to force generation is required for integrin activation and further maturation of NAs (66). In addition, our data reveals the existence of a large percentage of inactive α_5_β_1_ and α_v_β_3_ integrins inside FAs with little spatial overlap between integrins and main adaptors. As integrin activation is highly dynamic and transient, we propose that this pool of inactive integrin nanoclusters might be required to increase the probability of new engagements between integrins and their adaptors once the active integrin nanoclusters become disengaged, stabilizing as a whole the FAs for lifetimes much longer than the binding-unbinding dwell-times of integrins with their partners.

Quantification of super-resolution data showed a higher number of α_5_β_1_ nanoclusters compared to their α_v_β_3_ counterparts inside FAs. The relative percentage of active β_1_ nanoclusters is also higher than that of α_v_β_3_ and moreover these two integrin nanoclusters never intermix inside FAs. Interestingly, we found that while the nanocluster properties of α_5_β_1_ and α_v_β_3_ integrins in terms of number of localizations and size were independent of FA maturation, their spatial distribution inside FAs was markedly different as a function of seeding time. Indeed, α_5_β_1_ nanocluster distribution was already established at 90 min, while α_v_β_3_ nanocluster distribution appeared rather random at earlier seeding times and progressively organized reaching a well-defined lateral spacing at 24 hours of spreading time. As FA strengthening over time requires mechanical forces, our data strongly suggest that forces might play a differential role in the lateral distribution of both integrin nanoclusters over time. Further experiments lowering actomyosin contractility of the cells combined with the use of mechanical stretching devices or substrates of different stiffness would be required to further investigate this hypothesis.

Importantly, we found that active β_1_ nanoclusters preferentially locate at the periphery of FAs and in proximity with paxillin, talin and vinculin nanoclusters, while active β_3_ are more uniformly distributed inside FAs. The differential degree of α_5_β_1_ and α_v_β_3_ in terms of expression, activation state, and lateral nano- and meso-scale segregation, strongly indicates that integrin activation and their distinct functions inside FAs are spatially regulated (Fig. 6). Based on our results we postulate that *(i)* α_5_β_1_, through the ring on the FA periphery and its close proximity to paxillin, talin and vinculin nanoclusters, primarily functions for substrate attachment in its high tensional state, engaging with actomyosin filaments; *(ii)* α_v_β_3_ and α_5_β_1_ integrin nanoclusters in the central region of the FA undertake the role of mechanotransduction, being more dynamic in terms of activation and inactivation states and most probably engaged with shorter branched actin filaments; and *(iii)* the difference in these roles is controlled by the distinct mechanical forces exerted on the integrins, depending on their location inside FAs, and for α_5_β_1_ perhaps also on its different tensional states and/or degree of engagement with the actin cytoskeleton (67).

**Figure 6.**
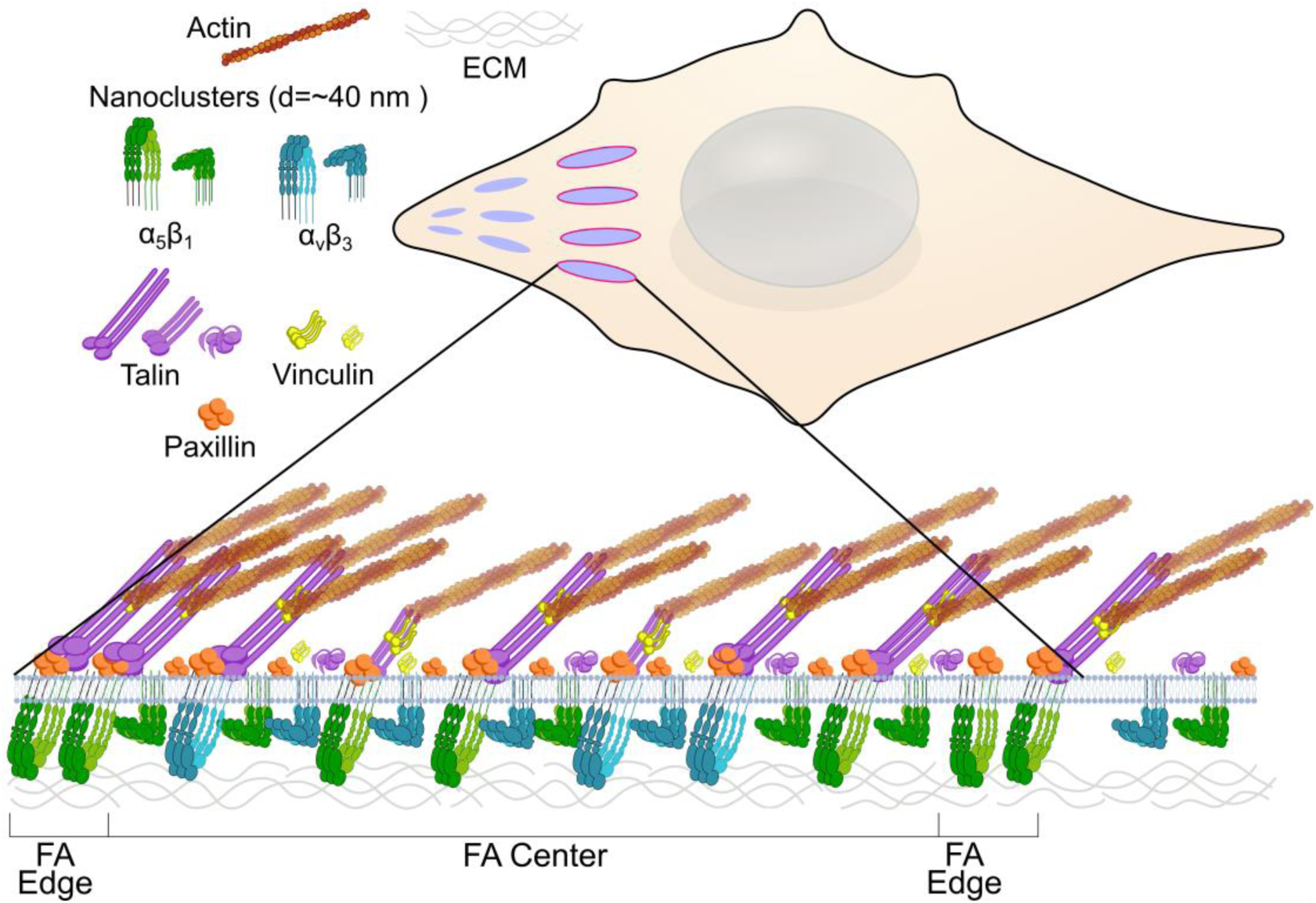
Spatial regulation and activation of integrin nanoclusters and protein adaptors inside focal adhesions. α_5_β_1,_ α_v_β_3_, and main protein adaptors organize as small nanoclusters inside FAs physically segregated from each other. A considerable high fraction of α_5_β_1_ and α_v_β_3_ integrins remain in an inactive state and do not intermix at the nanoscale. Active β_1_ nanoclusters (green) are preferentially enriched at the FA edges and reside in close proximity to the adaptor protein nanoclusters, possibly engaged to actomyosin filaments (brown). Integrin nanoclusters in the FA center are more randomly distributed with respect to one another, most probably engaged with branched cortical actin. According to our results we propose that α_5_β_1_ nanoclusters located at the FA periphery could be primarily involved in cell substrate attachment, whereas α_5_β_1_ and α_v_β_3_ nanoclusters at the center of FAs engage in mechanosensing and mechanotransduction. For simplification, we show a cross-section of a FA where active β_1_ integrins at the edge are engaged with talin. However, our data also show the presence of active β_1_ nanoclusters spatially separated from talin and possibly engaged to other binding partners such as tensin-3 linking β_1_ to the actin cytoskeleton (not shown in the cartoon for the sake of clarity).

Intriguingly, our analysis showed that active β_1_ nanoclusters preferentially locate at the FA edges regardless of their proximity to talin nanoclusters (Fig. 5B). These data are striking since active integrins are expected to be bound to both its ligand and talin, and therefore we had anticipated the whole active β_1_ population to be in close proximity to talin nanoclusters. We can envisage two plausible explanations for active β_1_ to be far from talin. One possible scenario is the nature of talin itself. As a long rod-like protein, talin stretching experiments have shown that the N- and C- termini can be separated up to 250 nm due to actomyosin activity (68). Since in our experiments, we have so far detected talin by labeling it in its C-terminal domain, which is opposite to the integrin binding side, it is possible that active β_1_ is engaged to extended talin even for inter-cluster distances larger than the median *i*NND. Further experiments labeling talin at the closest proximity to the integrin binding site could address the influence of talin size and extension in relation to β_1_ nanocluster spatial proximity.

A second possibility is the existence of other adaptor protein(s), aside from talin, able to activate β_1_. Such a candidate might be tensin-3, which has been shown to activate integrins (in particular, if β_1_ is phosphorylated) by interacting with the actin cytoskeleton, providing mechanical coupling for integrin activation (69) . In addition, it has been recently shown that tensin might compete with talin to bind α_5_β_1_ during the formation of fAs from FAs (70). Interestingly, our dual color STED experiments labeling the total α_5_β_1_ or active β_1_ populations together with tensin-3, reveal a clear co-enrichment of these proteins at the FA periphery (Fig. S8), suggesting that active β_1_ nanoclusters at the periphery of FAs might engage to the actin cytoskeleton either by associating to talin and vinculin or to tensin-3. Why would α_5_β_1_ be preferentially located at the FA periphery and associating either with talin or tensin? Although we do not know yet what drives the preferential location of α_5_β_1_ nanoclusters to the FA periphery, it is known that KANK2 also exhibits enrichment at the FA periphery, regulates talin activity and it is involved in the formation of α_5_β_1_-enriched fAs (71). Therefore, it is highly probable that α_5_β_1_ enrichment at the FA periphery is a necessary step for their translocation from mature FAs to fAs to then assemble fibronectin into the fibrillar networks as found and needed in connective tissues. Consistent with this idea, we have observed similar α_5_β_1_ distribution on mouse embryonic fibroblasts, which are the main cells that produce fibrillar adhesions. Thus, α_5_β_1_ nanocluster distribution inside FAs might be physiologically important for the process of fibronectin fibrillogenesis. This hypothesis is further substantiated by our observations of tensin-3 and β_1_ co-enrichment at the FA periphery as it has been documented that tensin is important for fibronectin fibrillogenesis (72) and more recently, it has been shown that tensin-3 interaction with talin drives the formation of fibronectin-associated fAs (70). As such, α_5_β_1_ enrichment at the FA periphery could serve to efficiently translocate these integrins from FAs to fAs, probably in a talin-tensin dependent manner. Additional experiments including simultaneous super-resolution mapping of α_5_β_1_, talin and tensin3 in mature FAs will be required to fully validate our hypothesis.

In summary, in contrast to the classical textbook view where FAs are pictured as dense and homogenous assemblies of integrins fully engaged to their cytoplasmic adaptors, our work reveals a highly complex organization inside FAs, where regulation of different integrins and their activation state are highly coordinated in space. We believe that such spatial regulation provides the means for different integrins to synergize and/or diversify their functions in terms of adhesion, migration, mechanosensing and mechanotransduction. Earlier reports showed that proteins such as KANK1, KANK2, and β_1_ integrins are enriched at the edges of adhesions, with KANK2 reducing the force exerted by the cells through its binding to talin (71, 73). KANK proteins have also been implicated in enabling communication between microtubules and actin by binding to talin and liprin-β_1_ at FA edges (74). More recent data suggest that the localization of KANK1 around FAs is driven by liquid-liquid phase separation (75). Our findings together with those mentioned above suggest that the α_5_β_1_ nanocluster belt is an important feature of FAs and may be critical for their function and/or the translocation of these integrins to fAs. However, further research is needed to fully understand how the organization of these key proteins contributes to their roles in fAs. p130Cas- and FAK-dependent phase separation has been also suggested to promote integrin clustering and NA assembly (76). Whether phase separation also contributes to the preferential organization and activation of β_1_ at the FA edges as we reveal here is an intriguing topic that deserves further investigation.

## Materials and Methods

### Sample preparation and immunolabeling

HFF-1 cells (human foreskin fibroblasts) were obtained from ATCC and used at low passage number (passage <15). Cells were maintained in DMEM supplemented with 15% FBS and cultured in a humidified atmosphere at 37° C and 5% CO2. Experiments were performed using cells seeded on 8-well chambered cover glass slides (#1 Lab-Tek, Thermofisher, #155411) previously coated with fibronectin (FN) at full substrate coverage. FN coating was performed by incubating the wells with 10 µg/ml FN for 30 min at room temperature (RT). We note that FBS can contain vitronectin (VN) and other proteins that may act as ligands of integrins. However, since the glass slides are saturated with FN prior to incubation with cells and medium, FN remains the dominant ligand in our experiments. After seeding, cells were allowed to spread for 24 h at 37° C and 5% CO2, after which they were fixed with 4% PFA in PBS for 15 min at RT. Cells were permeabilized using 0.1% Triton X-100 in PBS for 10 min and blocked with 3% BSA in PBS for 40 min. Details of reagents used for cell culture and sample preparation are provided in Table S1. For STORM experiments, primary antibodies against total α5β1 (CD49e/ IIA1) and αvβ3 integrins (CD51/61), active β1 integrin (CD29- 9Eg7) and active β3 integrins (CD61-LIBS2), and the adaptor proteins paxillin (Y113), talin1 (actin-binding C-terminal epitope) and vinculin were diluted in blocking buffer and added to the cells for 30 min at RT. Details of the primary antibodies used across the different experimental approaches are provided in Table S2. After washing with PBS, secondary antibodies that were conjugated in-house with dye pairs Alexa Fluor 405-Alexa Fluor 647 (405-647 channel) or Cy3B-Alexa Fluor 647 (Cy3-647 channel) (see Table S3), were added to the cells for 30 min at RT. After washing with PBS, cells were stored in PBS until imaged.

### Image acquisition

Single-molecule localization microscopy (SMLM) data were obtained on a commercial Nikon Eclipse Ti system (N-STORM), equipped with a 100x oil objective with NA 1.49 using TIRF illumination. The detector was an ANDOR technology EMCCD iXon 897 camera, with a 256x256 pixel ROI and a pixel size of 160 nm. The system has an Agilent technologies laser box with wavelengths of 405 nm, 560 nm and 647 nm. Details of the DNA-PAINT acquisition, performed on the same microscopy platform, are provided in the Supplementary Information section. For STORM experiments, imaging was performed in a buffer containing 100 mM of the reducing agent cysteamine mercaptoethylamine (MEA) (77 mg of MEA in 1 ml of 360 mM HCl, stock solution 1 M), the enzymatic oxygen scavenger Glox solution (0.5 mg/ml glucose oxidase, 40 μg/ml catalase), and 6.25% glucose in PBS. Details of the STORM imaging buffer reagents are found in Table S6. Dual color imaging was carried out cyclically and sequentially. In brief, we first detected the 405-647 channel in a 5-frame cycle consisting of 1 activator frame (405 nm laser on) followed by 4 reporter frames (647 nm laser on), followed by the Cy3-647 channel detection sequence, similarly composed of 1 activator frame (560 nm laser on) followed by 4 reporter frames (647 nm laser on). The frame rate was 50 Hz (20 ms per frame) for all frames. Imaging was carried out by cyclically repeating this 10-frame sequence until all the reporter’s dyes were exhausted (typically ∼70,000 frames).

### Single molecule detection, localization precision, and SMLM image reconstruction

Single-molecule localization microscopy images were reconstructed using Insight3 software, provided by Bo Huang (UCSF, initially developed in Xiaowei Zhuang’s lab, Harvard University). In each imaging frame, individual molecules were identified using intensity thresholding (gray levels set between 600-60,000), then each intensity profile was fitted with a 2D Gaussian function to identify the central position, i.e., the coordinates that represent the spatial localization of that fluorescent molecule. The analysis was run over the entirety of each data set, resulting in a final output that is a list of x,y coordinates (localizations) corresponding to each individual molecule that has been identified. Insight3 software allows for drift correction using the correlated function of the image, for details see (77). For Dual-color STORM, the main source of crosstalk is the nonspecific activation of the Alexa 647 by the 647 nm laser (78), which is activator-independent and can occur in any frame. We implemented statistical crosstalk correction where the Insight3 software analyses the full image in 50x50 nm regions of interest (ROIs) and determines if each ROI is predominantly one color of the other. Based on the acquisition design (in our case, one activator frame followed by four reporter frames), the software determines the probability of detecting a nonspecific reporter molecule (78). To minimize color crosstalk, only those localizations detected in the second frame of each five-frame STORM imaging cycle (that is, the first of the four reporter frames) were considered for future analytical steps. Since each single molecule can experience numerous blinking events, the variation of the x,y coordinates from frame to frame determines the effective resolution (79). To measure this, we detected multiple localization events arising from individual molecules deposited on glass and calculated the standard deviation of the localization positions. We measured the effective resolution of our system to be ∼15 nm. These localizations detected from nonspecific antibodies on glass were subsequently used to set a threshold to define the minimum number of localizations required for a cluster to be included in the DBSCAN-based nanocluster analysis. We found that the antibodies on glass had a median number of localizations of 5 (for channel 1) and 3 (for channel 2). With the aim of discarding all localizations coming from nonspecific binding or binding to individual antibody-labelled proteins, we set the threshold at 10 localizations per nanocluster.

### Generation of masks and classification of the localizations belonging to FAs, fAs or outside adhesions

A Fiji plugin, developed by the Lakadamyali lab, was implemented as an initial step to split the localization data from the STORM images into separate data sets corresponding to the localizations contained within defined ROIs (80). First, the data from the two channels of the reconstructed STORM image were imported and the localizations rendered together in a 1024x1024 pixels image (Fig. S1A). Next, we applied an intensity threshold, until the adhesion areas were visually detected, a Gaussian blurring of the image (by 2 pixels), and a second threshold filter until a smooth binary mask was obtained (Fig. S1B). The final mask retaining only the adhesion areas was obtained by manually selecting the regions corresponding to the adhesion complexes (identified by visual inspection based on their size and location). In the first steps, this Fiji plugin creates separate files containing the localizations that fall within the selected ROIs: all adhesions, glass, and outside adhesions (i.e., cell membrane that contain the remaining areas of the image that do not correspond to all adhesions or glass). In a second step the “all adhesions” file was again analyzed by the same plugin and all adhesions were split into FAs and fAs and rendered with Insight3. Fig. S1C-F shows the resulting rendered localizations for the different regions. These final two-color files for different cell regions were then individually and independently analyzed.

### Image analysis

#### Clustering detection using the DBSCAN algorithm

DBSCAN is a density-based data clustering algorithm designed to be executed on arbitrary data sets with minimal input from the user (40). The algorithm works by identifying arbitrary-shaped clusters of data points using two user-defined input values: epsilon (ε), which is a radial distance that serves to define a local neighborhood from a randomly selected initial data point; and Nmin, which is the minimum number of points required to be within the initial point’s neighborhood, so it is considered part of the cluster seed. For our image analysis, spots on glass outside the cell regions were analyzed using ε = 20 nm and Nmin = 2, in order to define a minimum threshold for the number of localizations per nanocluster. Important to note that we carried out this analysis on glass areas, next to the cells, found in all of our STORM images. In this way we ensure having the same imaging conditions for in-cell nanoclusters and for spots-on-glass, however, at the expense of having a broader distribution of localization per spot-on-glass due to the presence of cell debris on the glass. When analyzing the nanoclusters found in ROIs within the cell, the DBSCAN parameters were set to ε = 20 nm and Nmin=3 localizations, which are common values used in the field (36, 45). Nevertheless, since multiple blinks of a fluorophore might result in artifactual clusters, we only considered true nanoclusters as those containing 10 or more localizations. Examples on the identification of different nanoclusters by DBSCAN are shown in Fig. S1G. The algorithm also provides the center of mass (CoM) coordinates of each nanocluster together with the number of localizations per nanocluster and their diameter.

#### Nearest neighbor distance (NND) analysis

To establish how the protein nanoclusters are distributed relative to one another, we used the MATLAB (MathWorks) function *knnsearch*. This function computes for each nanocluster the NND, which is the shortest distance from the CoM of a given nanocluster to its closest CoM nanocluster neighbor. We also used the same function to determine the *i*NND (i.e., NND between nanoclusters of different proteins). To measure the percentage of nanoclusters that overlap (colocalization fraction), we first computed the edge-to-edge distance between nearest neighbors (EED-NN) by subtracting from the NND the radii of the two-corresponding near-neighbor nanoclusters. From the EED-NN distributions we calculated the percentage of occurrences with EED-NN values below zero. The *knnsearch* function was also used to extract the distance-to-the-FA edge, i.e., the shortest distance between nanoclusters and the edge of adhesions. To that end, we computed the NND between the CoM of each nanocluster to the center coordinates of the list of pixels that fall on the edge of the FA mask.

### Computational generation of uniform nanocluster distributions

To assess whether the distance distributions obtained from the experimental data (NND, EED-NN and distance-to-the-FA-edge), corresponded to a preferential type of organization or to a random spatial distribution, we performed simulations by generating in silico histograms of non-overlapping and randomly distributed nanoclusters inside FAs. For this, we first extracted the exact number and size of nanoclusters per adhesion from the experimental data. Our initial approach was to computationally generate the same number of nanoclusters and distribute them uniformly over the adhesion masks regardless of their physical size. However, this approach was not computationally suitable because, in many cases, large nanoclusters did not fit without overlap in the remaining space available within adhesions after randomly positioning the large majority of smaller nanoclusters. We therefore classified nanoclusters according to their physical size and positioned the largest nanoclusters first, followed by the remaining smaller ones. In addition, because our experimental data showed well-segregated nanoclusters, we imposed the condition that simulated nanoclusters should not overlap with one another. Fig. S9 shows the different steps followed to generate distributions of random nanoclusters inside FAs.

### Statistical analysis

Statistical analysis was carried out using GraphPad Prism (version 9.12). The details of the individual statistical tests as well as the level of significance along with p-values are explicitly provided in the figure legends. No predetermined statistical method was employed for establishing the sample size, as the sample sizes were not selected based on a predefined effect size. In this context, multiple experiments were independently performed, employing various sample replicates, as detailed in Table S8.

## Supplemental material

The supplemental material contains four supplementary texts associated with the full methodology used for STED imaging (Supplementary Text 1) and DNA-PAINT (Supplementary Text 2), description of the Voronoi-Density Island clustering algorithm for STORM data analysis (Supplementary Text 3) and description of the computational generation of uniform nanocluster distributions (Supplementary Text 4). Supplemental material also contains eight different Tables that specify the reagents used for cell culture and sample preparation, the primary antibodies used in the experiments, the secondary antibodies and their fluorophores used in STORM, DNA-PAINT and STED experiments, a list of the reagents and dyes used for STORM and DNA-PAINT imaging, and finally, a summary of experiments analyzed either as individual proteins or in protein pairs. In addition, the supplementary material contains nine different supplementary figures together with their captions. Fig. S1 provides information on the analysis pipeline used to identify and characterize nanoclusters of the different proteins. Fig. S2 provides quantification of the nanocluster sizes and number of localizations inside FAs and outside ACs for all the proteins investigated. Fig. S3 shows STED imaging of protein nanoclustering inside and outside FAs. Fig. S4 shows DNA-PAINT super-resolution images and nanocluster quantification inside and outside FAs. Fig. S5 provides quantification of the STORM data using the Voronoi-tessellation approach. Fig. S6 provides quantification of the edge-to-edge distance for the protein nanocluster pairs investigated and percentage of nanoclusters that physically overlap with each other. Fig. S7 provides quantification of how different protein nanoclusters are organized with respect to the FA edge as well as quantification of *i*NND distributions between active β_1_ nanoclusters and their adaptor partners paxillin, talin and vinculin. Fig. S8 shows dual color STED images and quantification of α_5_β_1_ and tensin on the FA periphery. Fig. S9 illustrates the different steps followed to generate in-silico distributions of non-overlapping, randomly distributed nanoclusters inside FAs.

## Data Availability Statement

The data supporting the findings of this study are available from the corresponding author upon reasonable request. The codes used in this study are available from the corresponding author upon reasonable request.

## Acknowledgments

The authors would like to thank J. Andilla and M. Rivas for technical support and to the Lakadamyali lab for the Fiji plugin used to split data into ROIs. The research leading to these results has received funding from the European Commission H2020 Program under grant agreement ERC Adv788546 (NANO-MEMEC) (to M.F.G.-P.), Government of Spain (Severo Ochoa CEX2019-000910-S, State Research Agency (AEI) (PID2020-113068RB-I00 / 10.13039/501100011033 and PID2023-147711NB-I00 (to M.F.G.-P.), RYC-2017-22227 and PID2022-138282NB-I00 project funded by the MCIN/AEI/10.13039/501100011033/ FEDER, UE (to F.C.)), Fundació CELLEX (Barcelona), Fundació Mir-Puig and the Generalitat de Catalunya through the CERCA program and AGAUR (Grant No. 2021 SGR01450 M.F.G.-P.). N.M. acknowledges funding from the European Union H2020 under the Marie Sklodowska-Curie grant 754558-PREBIST. The authors declare no competing financial interests.

## Author Contributions

S.K prepared the biological samples, performed STORM imaging and analyzed the data. A.G.-G. prepared the biological samples, performed STED and DNA-PAINT imaging and analyzed the data. N.M. developed algorithms for data analysis and contributed to the analysis. F.C. and M.F.G.-P. supervised the research. S.K., A.G.-G., F.C. and M.F.G.-P conceived and discussed the experiments, interpreted the data, and wrote the manuscript. All the authors read, provided feedback and approved the manuscript.

## Notes

### Competing Interest Statement

The authors have declared no competing interest.

### Summary of Updates

In the revised manuscript, we have updated the text in the first section of the results "Proteins of the ACs organize as universal nanoclusters across regions of the basal membrane" to include new control experiments using alternative super-resolution microscopy approaches STED (supplementary Text 1 and Fig S3) and DNA PAINT (supplementary text 2 and Fig S4). Additionally we reanalysed our STORM data using Voronoi tessellation as an alternative cluster identification approach and compared the new results to previously analysed images using DBSCAN (supplementary text 3 and Fig S5). We included a new results section "Lateral organization of integrin nanoclusters inside FAs" and Figure 2, to provide a first insight on the role of mechanical forces in the distribution of different integrin nanoclusters inside FAs. Here, we have performed experiments at different cell seeding times where it is known that FA maturation over time requires mechanical forces. Finally, we provide dual color STED images of integrin b1 together with tensin showing co-enrichement of both proteins at the FA periphery, suggesting that b1 at the FA edges might be engaged with talin or tensin for b1 integrin activation and engagement with the actin cytoskeleton (Fig. S8). We have updated our model (Figure 6) and extended our discussion to incorporate our new findings. The author list has been updated to include Amaris Guevara-Garcia as second author for her contributions to the paper in the form of extensive control experiments and dual color STED imaging and analysis. The supplementary information has been updated to include all the new methods and their results (supplementary text 1-3 and Fig S3-5). Additional Tables in the supplementary information provide details on the reagents used in the control experiments.

