## Supplemental Text and Figures for "Differential spatial regulation and activation of integrin nanoclusters inside focal adhesions"

### Supplementary information

### Supplementary Text 1: Methodology for STED imaging

*Sample preparation and immunolabeling*

For STED experiments, sample preparation, fixation, permeabilization and blocking were performed as described in the main Materials and Methods section. Cells were labeled with primary antibodies against total $\alpha_5\beta_1$ integrin (CD49e/ IIA1), active $\beta_1$ integrin (CD29- 9Eg7), total $\alpha_v\beta_3$ integrin (CD51/61), paxillin (177) or Tensin3, (Table S2), cells were washed with PBS and incubated for 60 min at RT with secondary antibodies conjugated to Abberior STAR dyes, Abberior GmbH (Table S4) diluted in blocking buffer. After washing with PBS, cells were stored in PBS until imaged.

*Image acquisition*

STED imaging was performed on an Abberior INFINITY platform built around an Olympus IX83 inverted microscope, equipped with a 100× oil-immersion objective (NA 1.4). For STAR RED imaging, excitation was performed using the 640 nm pulsed laser (10% power) with depletion provided by the 775 nm pulsed STED laser (50% power). For STAR ORANGE imaging, excitation was performed using the 561 nm pulsed laser (15% power) with depletion provided by the 775 nm pulsed STED laser (40% power). Images were acquired with 15-line accumulation. Acquisition parameters were kept constant within each experimental condition.

*Image analysis*

Single-color STED: Line intensity profiles were extracted from raw STED images using Fiji/ImageJ by manually drawing line ROIs across focal adhesion regions.

Dual-color STED: Dual-color images were quantified using a custom Fiji/ImageJ–MATLAB pipeline. For each FA, radial mean intensity profiles were computed inward from the FA edge in radial bins, restricted to distances ≤600 nm from the FA boundary. Profiles were background-corrected by subtracting the minimum intensity value and then normalized to the range [0–1] to reduce variability in absolute signal levels and emphasize relative spatial distributions. Profiles were then interpolated onto a common radial grid and averaged across FAs, with variability reported as the standard error of the mean (SEM). Edge enrichment was quantified by comparing mean normalized intensities in the edge region (60–180 nm from the FA boundary) to those in the interior (180–600 nm), calculated as: Mean Intensity (Edge) / Mean Intensity (Interior). Values >1 indicate enrichment at the FA periphery. Statistical comparisons were performed using two-sided Welch's t-test

### Supplementary Text 2: Methodology for DNA-PAINT imaging

*Sample preparation and immunolabeling*

Sample preparation and primary antibody labeling were performed as described in the main Materials and Methods section. For DNA-PAINT experiments, cells were labeled with primary antibodies against active β1 integrin (CD29-12G10), αvβ3 integrin (CD51/61), paxillin (177) and talin1 (TA205/N-terminal epitope), as listed in Table S2. Following primary antibody incubation and washing with PBS, secondary labeling was performed using DNA-conjugated camelid single-domain antibodies (sdAbs; Massive Photonics DNA-PAINT; Table S5), following the manufacturer's instructions. The sdAbs were site-specifically coupled to a single DNA docking strand, with up to two sdAbs binding per primary antibody, resulting in a maximum of two docking sites per primary antibody. The small size of sdAbs minimizes linkage error and improves epitope accessibility, reducing spatial uncertainty in localization-based measurements. After labeling, cells were washed with PBS and stored until imaging.

*Image acquisition*

DNA-PAINT imaging was performed on the same Nikon Eclipse Ti N-STORM microscope described in the main Materials and Methods section. The detector was a Hamamatsu ORCA-Flash4.0 CMOS camera. Samples were imaged in Massive Photonics DNA-PAINT imaging buffer containing 1.5 nM imager strands complementary to the docking strands (see Table S7). Excitation was provided by the 647 nm laser, and images were acquired with a 100 ms exposure time for a total of 20,000 frames.

*Image reconstruction and nanocluster analysis*

DNA-PAINT image reconstruction and nanocluster analysis followed the same pipeline described in the main Materials and Methods for STORM data, including single-molecule detection and localization using Insight3, drift correction, FA segmentation using the Fiji plugin developed by the Lakadamyali laboratory.

For DBSCAN-based nanocluster identification, isolated anti-mouse sdAb–ATTO655 spots on glass, located in proximity to cellular regions, were used to estimate the number of localizations corresponding to individual binding sites under the same imaging conditions, as in the STORM analysis. The mean number of localizations per individual spot-on glass was 5.43 ± 1.18. Based on this value, DBSCAN-based nanocluster identification was performed using ε = 20 nm and MinPts = 10 localizations.

### Supplementary Text 3: Hybrid Voronoi–Density Island Clustering of STORM *Localizations*

Clustering of single-molecule localizations was performed using a two-step procedure that combines first-rank Voronoi tessellation with a density-island segmentation approach. This hybrid strategy is adapted from previous work (1-4) and incorporates modifications to enhance robustness across heterogeneous densities and complex nanostructures.

To quantify local localization density, we computed the first-rank Voronoi density for each localization. This metric extends the classical Voronoi density by incorporating contributions from immediate Voronoi neighbors and has been shown to improve stability in regions with strong local density gradients. For a localization *i*, the first-rank density is defined as:

$$\delta_i^{1st} = \frac{1 + \sum_j neighbors_j}{A_i + \sum_j A_j}$$

where the numerator counts the localization itself plus all Voronoi neighbors that share a cell boundary with it, and the denominator is the sum of the corresponding Voronoi areas. We normalize this quantity by the global density of a random point cloud,

$$\widehat{\delta_i^{1st}} = \frac{\delta_i^{1st}}{\delta_{rand}}$$

where the density of random localizations is computed as the total number of localizations divided by the total area of the region of interest (ROI). The ROI is the minimal bounding box containing all experimental localizations.

Rather than adopting a fixed zero-threshold as in (1), we first generated a random point distribution of equal size, computed its normalized first-rank Voronoi densities, and evaluated their cumulative distribution function (CDF). A user-defined percentile (cdf_threshold) was applied to this distribution to obtain an adaptive cutoff. Experimental localizations with normalized first-rank density above this value were classified as high-density and used as inputs for the clustering stage. Lower CDF thresholds yield more inclusive sets of high-density points, whereas higher thresholds restrict analysis to the most densely packed regions. A threshold value was set to 70 for all data sets.

Second, we performed a density-island segmentation and identification of cluster centers. High-density localizations were converted into 12 nm–pixel images in which each pixel intensity corresponded to the number of localizations it contained. These images were smoothed using a 5×5 pixel convolution kernel to reduce noise and highlight coherent density regions.

A fixed intensity threshold was then applied to generate density islands, defined as connected regions of above-threshold signal. Each island represents a spatially continuous aggregation of high-density points and may contain one or multiple molecular clusters.

To identify cluster centers within each island, local maxima were detected in the smoothed density image. Since the MATLAB version used in this study predates the islocalmax function, we implemented a custom two-dimensional maxima detection routine (islocalmax2_custom.m) providing equivalent functionality. For islands containing one density maximum, the cluster centroid was assigned as the center of mass (CoM) of all localizations in that island.

For islands containing multiple maxima, localizations were initially assigned to their nearest maximum using a knnsearch-based nearest-neighbor classification. A provisional CoM was computed for each group, and a second reassignment was performed to refine cluster membership. This iterative refinement compensates for the fact that maxima are initially extracted from a smoothed image, whereas final centroid positions must reflect the distribution of raw localizations.

Clusters containing fewer than a minimum number of localizations were excluded. For the remaining clusters, a polygonal hull was constructed from their assigned localizations to estimate cluster area.

Because the density image oversamples the underlying point cloud, heterogeneous or elongated structures may be artificially split during density segmentation. To mitigate this, clusters with centroids separated by less than 35 nm—a distance on the order of the STORM localization precision (~30 nm)—were merged, preventing over-segmentation of single clusters.

The MATLAB codes used for the analysis are available at
https://github.com/nmateosHub/VoroIslands_Clustering


**Supplementary Text 4:**

**Computational generation of uniform nanocluster distributions**

To assess whether the distance distributions (NND, EED-NN, d2e) obtained from the experimental data correspond to a preferential type of organization or to a random distribution, we performed simulations by generating in silico distance histograms of randomly distributed nanoclusters (according to a uniform distribution) inside individual adhesion structures. Our pipeline essentially needs two ingredients: *First*, to extract from the experimental data the exact number (and size) of nanoclusters per adhesion; and *second*, to distribute the same number of nanoclusters in a random fashion within the adhesions, while excluding their spatial overlapping. We therefore followed the different steps:

***Step 1:** Identification of nanoclusters inside a given adhesion.* We first paired the mask of the adhesions in a cell to the corresponding experimental data. Each particular adhesion, $A_n$, contains a given number of nanoclusters. The algorithm, coded in MATLAB, goes sequentially through each mask, identifies which adhesions contained nanoclusters, counts the number of nanoclusters belonging to each adhesion, extracts their center of mass (CoM) position, and measures their physical size.

***Step 2:** Sorting the nanoclusters as function of their physical size*. Although our initial approach was to computationally generate the same number of nanoclusters and distribute them uniformly over the adhesion masks regardless of their physical size, we found that this approach was not computationally suitable. The reason for it is that in many cases the large nanoclusters did not fit (without overlapping) in the remaining space available within the adhesions after randomly positioning the large majority of smaller nanoclusters. We thus opted for classifying the nanoclusters according to their physical size and then start our procedure by first positioning the largest nanoclusters and later all the remaining smaller ones. In addition, as our experimental data renders well segregated nanoclusters, we imposed in our algorithm the condition that nanoclusters should not overlap with one another.

To classify the nanoclusters as a function of size, we plotted the experimental nanocluster radii as a cumulative distribution function, determined a threshold nanocluster radius, $R_{th}$, which corresponds to the lower bound of the 80th percentile. We sorted the radii according to their size and segregated the nanoclusters into two groups: the 20% largest and 80% smallest (Fig. S9A). Therefore, the number of large clusters, $N_{big}$, for any adhesion was equal to the number of clusters N with a radius greater than the threshold radius, $r > R_{th}$.

***Step 3:** Random distribution of the largest clusters and size assignment.* First, we generated 100 x $N_{big}$ random points uniformly distributed over the smallest non-tilted boundary box containing the adhesion (Fig. S9B, *left panel*). We identified and kept only those points inside the adhesion (Fig. S9B, *middle panel*). If the number of points that fell inside the adhesion, $N_{in}$, was larger than $N_{big}$ we randomly removed points to ensure that $N_{in} = N_{big}$. If $N_{in} < N_{big}$ we then repeated the procedure to generate new sets of $N_{in}$ until $N_{in} = N_{big}$, (Fig. S9B, *right panel*). With $N_{in} = N_{big}$ achieved, we then randomly assigned a radius to each point in $N_{in}$ from the set of large radii determined by $r > R_{th}$.

***Step 4:** Identification and rejection of overlapping nanoclusters.* To identify nanoclusters that spatially overlapped, we calculated the center-to-center distance from each nanocluster to its *k*-th nearest neighbors, *k*ND, where if the number of nanoclusters in an adhesion was greater than 10, $N_{in} > 10$, then $k = 10$, and if $N_{in} \leq 10$ then $k = N_{in}$. We then calculated the edge-to-edge distances between nearest nanoclusters (i.e., EED-NN), which we defined as:

$$EED\text{-}NN = kND - R_{ref} - R_k, \quad \text{(Eq. S1)}$$

where $R_{ref}$ and $R_k$ correspond to the radius of the reference and the *k*-neighbor nanoclusters, respectively. If nanoclusters overlap, i.e., EED-NN $\leq 0$, for any of the *k*-neighbors, then both the reference nanocluster and the neighbor are rejected.

We followed this approach for all large nanoclusters, resulting in two possible outputs: nanoclusters that survived the non-overlapping criterion (Fig. S9C, *green dots*) and those rejected due to overlapping with a neighbor (Fig. S9C, *red dots*). Thus:

$$EED\text{-}NN > 0 \rightarrow XY_S, \text{with radius}, R_S \quad \text{(Eq. S2)}$$

$$EED\text{-}NN \leq 0 \rightarrow XY_{rej}, \text{with radius}, R_{rej} \quad \text{(Eq. S3)}$$

The number of nanoclusters that survived the non-overlapping criterion, $N_S$, were then fixed in place with their randomly assigned $XY_S$ coordinates and radius $R_S$. All the other nanoclusters that failed the non-overlapping criterion were rejected, $N_{rej}$.

***Step 5:** Generation of newly proposed random nanoclusters, $N_{prop}$*. We included new random points ($N_{prop}$ = $N_{rej}$) and assigned to each of them a radius from the set of radii initially rejected, $R_{rej}$ (see different panels in Fig. S9C).

***Step 6:** Assessment of nanocluster overlap between newly proposed nanoclusters and Ns.* The total number of randomly generated clusters was:

$$N_{tot} = N_S + N_{prop} \quad \text{(Eq. S4)}$$

We calculated the *k*ND as above, but this time we calculated them from each point in $N_{prop}$ to each point in $N_{tot}$. We once again calculated the EED-NN using Eq. S1, however, for this step if EED-NN ≤ 0 we only rejected the reference nanocluster from the newly proposed set. If EED-NN > 0, we added the newly proposed set to the $N_S$ pool. This step once again gave two outputs: an updated set of nanoclusters that survived along with the radii they were assigned, and a number of clusters that were rejected. We repeated the last two steps, generating newly proposed points equal in number to the previously rejected ones and assessing if they overlap with the pre-existing nanoclusters, until we achieved $N_{rej}$ = 0, and thus $N_S$ = $N_{big}$. In this way we generated a set of randomly distributed large nanoclusters that do not physically overlap with one another, and with a size that was randomly extracted from the experimentally measured nanoclusters that had r > $R_{th}$.

***Step 7:** Generation and placement of small nanoclusters, $N_{small}$.* With all large nanoclusters in place, we again repeated steps 5, and 6, using this time $N_{small}$, i.e., the remaining 80% of nanoclusters whose r < $R_{th}$. We generated newly proposed nanoclusters, with $N_{prop}$ = $N_{small}$ and evaluated if the newly proposed nanoclusters overlapped, either with themselves or with the larger nanoclusters placed previously. We did this sequential loop of generation and assessment until the number of small nanoclusters that were rejected due to overlap was less than or equal to 10, i.e., $N_{rej}$ ≤ 10.

***Step 8:** Placement of the last 10 random nanoclusters in the adhesion*. Once $N_{rej}$ ≤ 10, we placed the nanoclusters individually, one by one. To this end, we generated 1000 points inside the adhesion of interest and assigned them all the same radius, *r*, from the list of $R_{rej}$. We then assessed each of these 1000 nanoclusters and found those that had an EED-NN > 0, implying they did not overlap with another nanocluster. We then randomly selected one of these nanoclusters and added it to the survivor list with its radius, *r*, and *XY* coordinates. We repeated this cycle until $N_{rej}$ =0. We followed all these steps on all adhesions, and 10 times per cell to reach a close to theoretical random distribution of non-overlapping nanoclusters.

**Supplementary Tables**

Table S1: **Reagents for cell culture and sample preparation**

| Reagent | Abbreviation | Company | Cat # |
|---|---|---|---|
| **Fetal Bovine Serum** | FBS | Biowest | S181B |
| **Dulbecco's Modified Eagle Medium** | DMEM | Capricorn–Scientific | DMEM-HXRXA |
| **Trypsin** | | Biowest | L0910-100 |
| **Bovine Serum Albumin** | BSA | Capricorn–Scientific | BSA-1S |
| **Phosphate-Buffered Saline** | PBS | Capricorn-Scientific | PBS 1A |
| **Fibronectin** | FN | Sigma Aldrich | 11080938001 |
| **Paraformaldehyde** | PFA | Sigma Aldrich | F1635-500 |
| **Triton X-100** | | Fisher Scientific | BP151 |

Table S2: **Primary antibodies used in the experiments**

| Target Protein | Host | Stock conc* (mg/ml) | Working dilution | Company | Cat # | Comments |
|---|---|---|---|---|---|---|
| **$\alpha_5\beta_1$** | Ms | 0.5 | 1:100 | BD Biosciences | 555510 | CD49e/ IIA1 -extracellular |
| **$\alpha_5\beta_1$** | Rb | 1 | 1:500 | Sigma Aldrich | AB1928 | CD49e/polyclonal - intracellular |
| **Act $\beta_1$** | Rat | 0.5 | 1:200 | BD Biosciences | 553715 | CD29- 9Eg7 |
| **Act β1** | Ms | 1 | 1:400 | Abcam | Ab30394 | CD29-12G10 |
| **$\alpha_v\beta_3$** | Ms | 0.1 | 1:20 | Abcam | Ab7166 | CD51/61 - αV + β3 |
| **Act $\beta_3$** | Ms | 0.5 | 1:100 | Sigma Aldrich | MABT27 | CD61-LIBS2 |
| **Paxillin** | Rb | 0.14 | 1:100 | Abcam | Ab32084 | Y113/paxillin |
| **Paxillin** | Ms | 0.25 | 1:100 | BD Biosciences | 610568 | 177/paxillin |
| **Talin1** | Rb | 1 | 1:100 | Abcam | Ab71333 | Polyclonal/ C-Term. |
| **Talin1** | Ms | 1 | 1:100 | Merk | MAB1676 | TA205/ N-Term. |
| **Vinculin** | Rb | 1.0–1.3 | 1:100 | Sigma Aldrich | v4139 | Polyclonal |
| **Tensin3** | Rb | 0.3 | 1:100 | Sigma Aldrich | HPA055338 | Polyclonal |

*conc=concentration; Ms=Mouse; Rb=Rabbit

Table S3: **Secondary antibodies and their fluorophores used in STORM experiments**

| Target species | In-house conjugated dye pair activator | In-house conjugated dye pair reporter | Host | Stock conc* (mg/ml) | Working dilution | Company | Cat # |
|---|---|---|---|---|---|---|---|
| | (dyes per antibody) | | | | | | |
| **Mouse** | Alexa Fluor 405 | Alexa Fluor 647 | Donkey | 0.1 | 1:20 | Jackson Immuno Research | 715-005-150 |
| | (6:1) | | | | | | |
| **Rabbit** | Alexa Fluor 405 | Alexa Fluor 647 | Donkey | 0.1 | 1:20 | Jackson Immuno Research | 711-005-152 |
| | (6:1) | | | | | | |
| **Rat** | Alexa Fluor 405 | Alexa Fluor 647 | Donkey | 0.1 | 1:20 | Jackson Immuno Research | 712-005-150 |
| | (5:1) | | | | | | |
| **Mouse** | Cy3 | Alexa Fluor 647 | Donkey | 0.1 | 1:20 | Jackson Immuno Research | 715-005-150 |
| | (6:1) | | | | | | |
| **Rabbit** | Cy3 | Alexa Fluor 647 | Donkey | 0.1 | 1:20 | Jackson Immuno Research | 711-005-152 |
| | (6:1) | | | | | | |

*conc=concentration

Table S4: **Secondary antibodies and their fluorophores used in STED experiments**

| Target Species | Conjugated Fluorophore | Host | Stock conc* (mg/ml) | Working dilution | Company | Cat # |
|---|---|---|---|---|---|---|
| **Mouse** | STAR RED | Goat | 1 | 1:250 | Abberior | STRED-1001-500UG |
| **Rabbit** | STAR RED | Goat | 1 | 1:250 | Abberior | STRED-1002-500 UG |
| **Rabbit** | STAR ORANGE | Goat | 1 | 1:250 | Abberior | STORANGE-1002-500UG |
| **Rat** | STAR RED | Goat | 1 | 1:250 | Abberior | STRED-1007-500UG |

*conc= concentration

Table S5: **Secondary antibody used in DNA PAINT experiments**

| Target Species | Comment | Host | Stock conc* (mg/ml) | Working dilution | Company | Cat # |
|---|---|---|---|---|---|---|
| **Mouse** | SdAB-1A23 | Camelid | 5uM | 1:300 | Massive Photonics | n.a. |

*conc= concentration

Table S6: **Reagents and dyes for STORM imaging**

| Reagent | Abbreviation | Company | Cat # |
|---|---|---|---|
| **Glucose Oxidase (from Aspergillus niger)** | | Sigma-Aldrich | G2133 |
| **Hydrochloric-Acid** | HCl | Sigma-Aldrich | 258148 |
| **Cysteamine mercaptoethylamine** | Cysteamine MEA | Sigma-Aldrich | 30070 |
| **α-D-glucose** | | Sigma-Aldrich | 158968 |
| **Catalase from bovine liver** | | Sigma-Aldrich | C100 |
| **Cy3** | | Sigma-Aldrich | GEPA23001 |
| **Alexa Fluor 647 carboxylic acid succinimidyl (NHS) ester** | Alexa647 | Invitrogen | A20006 |
| **Alexa Fluor 405 carboxylic acid succinimidyl (NHS) ester** | Alexa405 | Invitrogen | A30000 |

Table S7: **Reagents for DNA-PAINT imaging**

| Reagent | Comment | Company | Cat # |
|---|---|---|---|
| **Imager- ATTO 655** | 1 µM in TE buffer (10 mM Tris, 1 mM EDTA, pH 8) | Massive Photonics | n.a |
| **Antibody incubation Buffer** | Commercial buffer; composition not disclosed by the manufacturer | Massive Photonics | n.a |
| **Washing Buffer** | Commercial buffer; composition not disclosed by the manufacture | Massive Photonics | n.a |
| **Imaging Buffer** | Commercial buffer; composition not disclosed by the manufacturer | Massive Photonics | n.a |

Table S8: **Summary of STORM experiments analyzed either as individual proteins or in protein pairs.**

| Protein/protein pairs | # Cells | # Samples | #Cells per Sample |
|---|---|---|---|
| $\alpha_5\beta_1$ | 41 | 14 | 4,4,2,3,1,2,2,2,3,4,4,4,2,4 |
| $\alpha_v\beta_3$ | 47 | 13 | 3,4,1,2,5,3,4,1,2,3,3,3,4,1,5 |
| Act $\beta_1$ | 28 | 15 | 3,3,2,3,2,6,2,3,4,2,3,4,4,1,5 |
| Act $\beta_3$ | 17 | 8 | 4,3,4,3,4,4,2,4 |
| Paxillin | 43 | 13 | 4,4,3,4,1,3,3,2,3,2,4,3,4 |
| Talin | 23 | 7 | 2,3,2,5,6,2,3 |
| Vinculin | 18 | 9 | 1,2,2,2,3,3,3,4,3 |
| $\alpha_5\beta_1$ + Paxillin | 8 | 2 | 4,4 |

| | | | |
|---|---|---|---|
| $\alpha_5\beta_1$ + Talin | 5 | 2 | 2,3 |
| $\alpha_5\beta_1$ + Vinculin | 5 | 3 | 1,2,2 |
| $\alpha_v\beta_3$ + Paxillin | 8 | 3 | 3,4,1 |
| $\alpha_v\beta_3$ + Talin | 7 | 2 | 2,5 |
| $\alpha_v\beta_3$ + Vinculin | 6 | 3 | 3,4,1 |
| $\alpha_v\beta_3$ + $\alpha_5\beta_1$ | 8 | 3 | 2,3,3 |
| Act $\beta_1$ + Paxillin | 13 | 5 | 3,3,2,3,2 |
| Act $\beta_1$ + Talin | 8 | 2 | 6,2 |
| Act $\beta_1$ + Vinculin | 7 | 2 | 3,4 |
| Act $\beta_3$ + Paxillin | 14 | 3 | 4,3,4 |
| Act $\beta_3$ + Talin | 3 | 1 | 3 |

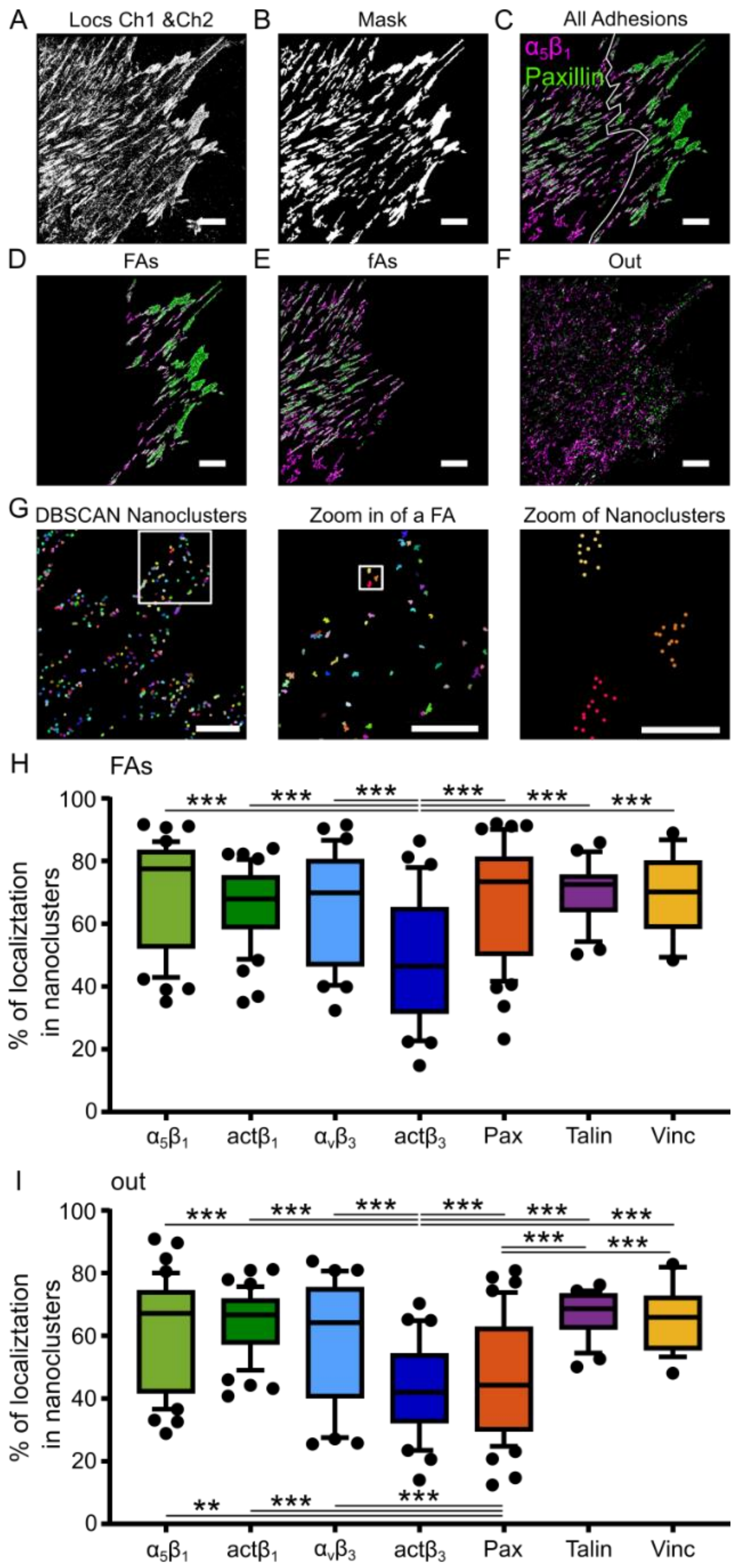
A
Locs Ch1 &Ch2
B
Mask
C
All Adhesions
$\alpha_5\beta_1$
Paxillin
D
FAs
E
fAs
F
Out
G
DBSCAN Nanoclusters
Zoom in of a FA
Zoom of Nanoclusters
H
FAs
% of localiztation in nanoclusters
$\alpha_5\beta_1$
act$\beta_1$
$\alpha_v\beta_3$
act$\beta_3$
Pax
Talin
Vinc
I
out
% of localiztation in nanoclusters
$\alpha_5\beta_1$
act$\beta_1$
$\alpha_v\beta_3$
act$\beta_3$
Pax
Talin
Vinc

**Figure S1: Steps of the analysis pipeline leading to nanocluster identification and characterization.**

*(A-G) Steps involved for analyzing the STORM data. (A) Image showing all the STORM localizations identified in channel 1 and channel 2. (B) Mask of all the ACs. (C-E) Insight3-generated rendering of the localizations identified to belong to ACs (C), where the white line shows the manually-drawn boundary line used to split ACs into FAs (D) and fAs (E). (F) Rendering of localizations found on the membrane outside (Out) the AC mask. (A-F) Scale bar: 5 µm. (G) The nanoclusters identified in a single channel by the DBSCAN algorithm (left), with a zoom-in on a FA (middle) and a further zoom into individual nanoclusters (right). Scale bar: 2 µm (left), 1 µm (middle) and 250nm (right). (H, I) Box-and-whisker plot showing the percentage of localizations found in nanoclusters in FAs (H) and on the membrane outside adhesions (I). The box represents the range from the 25$^{th}$ to the 75$^{th}$ percentiles and whiskers from 10$^{th}$ to 90$^{th}$ percentiles, with the median indicated by a horizontal line. Points outside these whiskers correspond to outliers. Individual points (points within the whiskers not shown) correspond to the median value over all nanoclusters for each cell (for N values, i.e., number of independent experiments, see SI Table S5). One-way ANOVA test, where only those pairs that were found to be significantly different are marked on the graph. ns = not significant, $p>0.05$; *, $p<0.05$; **, $p<0.01$; ***, $p<0.001$.*

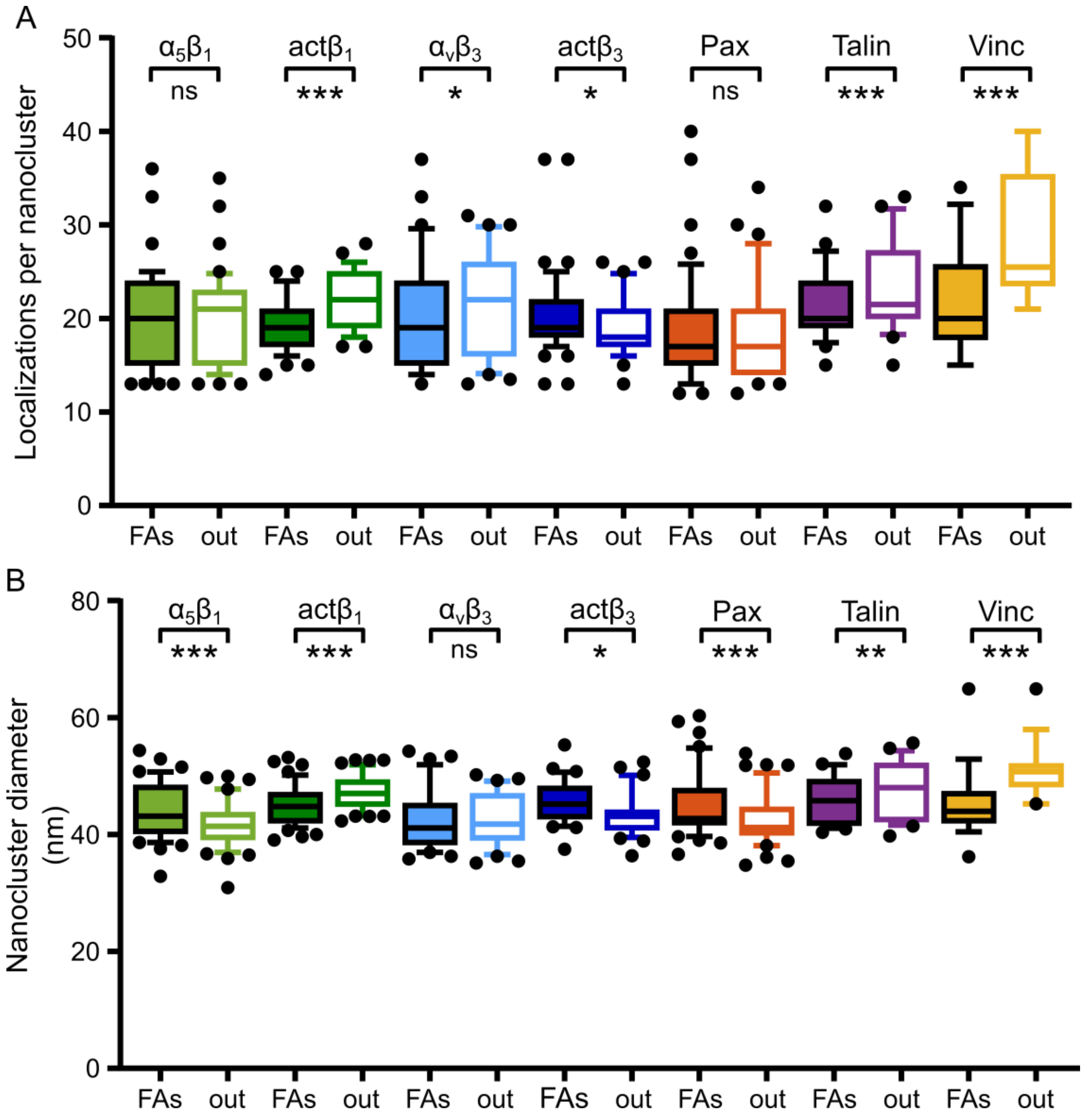


**Figure S2: Nanocluster characteristics inside FAs and outside ACs.**

*(A, B) Box-and-whisker plots showing the number of localizations per nanocluster (A) and the nanocluster diameter (B) for nanoclusters of each protein found inside FAs and outside ACs (out). The box represents the range from the 25$^{th}$ to the 75$^{th}$ percentiles and whiskers from 10$^{th}$ to 90$^{th}$ percentiles, with the median indicated by a horizontal line. Each data point corresponds to the median value over all nanoclusters for each individual cell. Points outside these whiskers correspond to outliers. Individual points (points within the whiskers not shown) correspond to the median value over all nanoclusters for each cell (for N values, i.e., number of independent experiments, see SI Table S5). Statistical differences were computed using a paired t-test between FAs and out for each protein, ns = not significant, p>0.05; *, p<0.05; **, p<0.01; ***, p<0.001.*

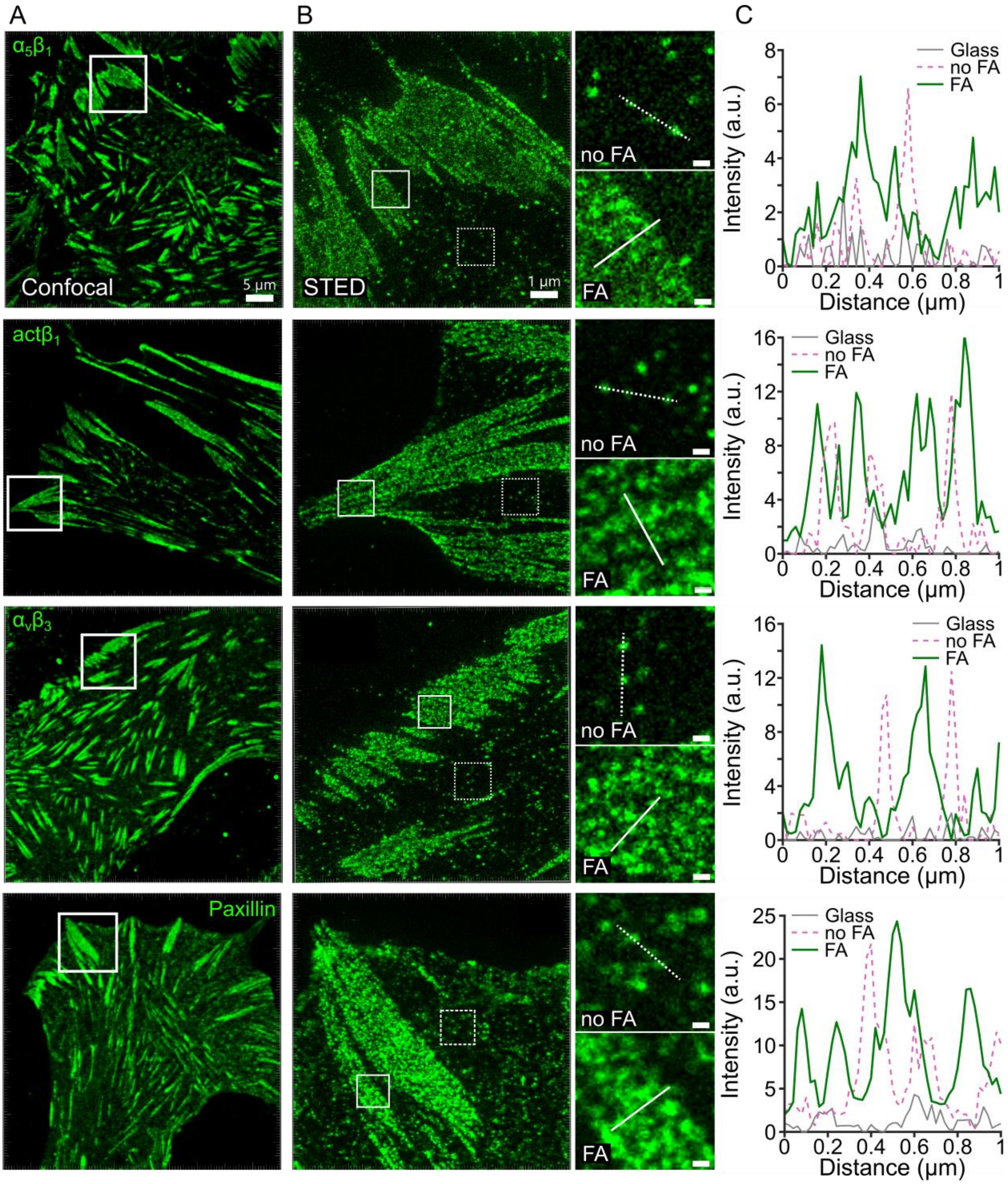


**Figure S3: STED imaging confirms nanoscale protein clustering inside and outside FAs.**
*(A) Confocal overview images of cells showing, from top to bottom, total $\alpha_5\beta_1$ integrin, active $\beta_1$ integrin, $\alpha_v\beta_3$ integrin, and paxillin. The white square indicates the focal adhesion (FA) region imaged by STED microscopy. Scale bar: 5 µm. (B) STED images of the indicated FA region. White squares indicate regions selected inside FA and outside FA for higher-magnification analysis. Insets show zoomed-in STED views highlighting fluorescence spots within regions inside FA, indicated by solid white squares, and outside FA (no FA) indicated by dashed white squares. Scale bars: 1 µm for the main images and 0.2 µm for the insets.* **(***C) Representative line intensity profiles extracted from STED images across individual fluorescence spots on glass, in no-FA regions, and in FA regions. Higher*

*intensities in FA and no-FA regions compared with individual fluorescence spots on glass support the presence of protein nanoclusters in both regions. Comparable intensity profiles inside and outside FAs suggest similar relative protein levels per nanocluster. Given the effective resolution of STED microscopy in our experiments (~ 60 nm), quantitative nanocluster analysis was performed using DNA-PAINT (see Fig. S4).*

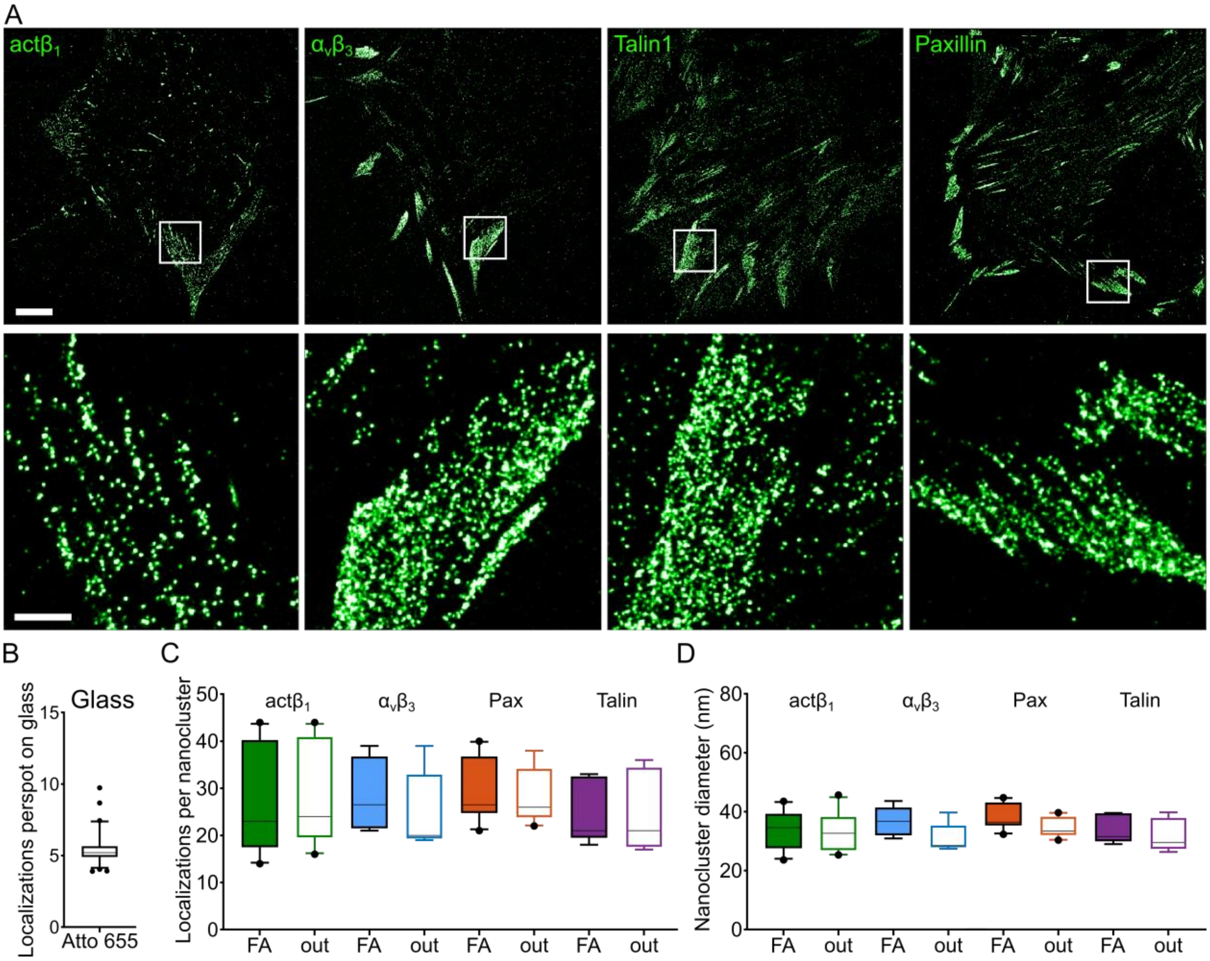


**Figure S4: DNA-PAINT confirms a conserved nanocluster organization of integrins and main adaptors across FA proteins.**

*(A) Top: Representative single-color DNA-PAINT super-resolution reconstructions of FA proteins labeled with anti-mouse sdAbs conjugated to ATTO655. Shown are (from left to right) active β1 integrin, αvβ3 integrin, talin1 and paxillin. Scale bar: 5 µm. Bottom: Magnified regions indicated in each panel. Scale bar: 1 µm. (B) Distribution of localization counts per binding site measured for isolated anti-mouse sdAb–ATTO655 spots immobilized on glass in proximity to cellular regions. The mean number of localizations per individual spot-on glass was 5.22 ± 1.18 . (C) Box-and-whisker plots showing the median number of localizations per nanocluster inside and outside FAs. (D) Nanocluster diameter for the investigated proteins. Each data point corresponds to the median value of one cell containing multiple focal adhesions (n = 10 cells per condition).*

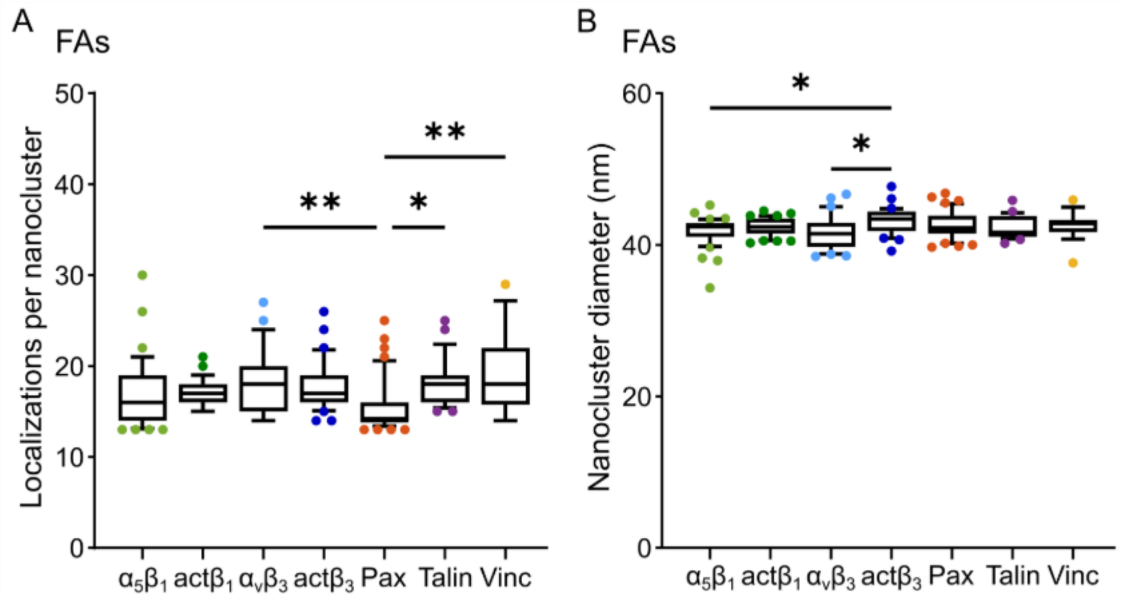


**Figure S5: Analysis of the STORM data using Voronoi-tessellation:**
*Box-and-whisker plots showing the number of localizations per nanocluster (A) and the nanocluster diameter (B) for the proteins in this study. The box represents the range from the 25th to the 75th percentiles and whiskers from 10th to 90th percentiles, with the median indicated by a horizontal line. Points outside these whiskers correspond to outliers. Individual points (points within the whiskers not shown) correspond to the median value over all nanoclusters for each cell (for N values, i.e., number of independent experiments, see SI Table S8). Statistical differences were found using the multi comparison one-way ANOVA test, P: 0.12 (ns), 0.033(*), 0.002 (**), <0.001(***).*

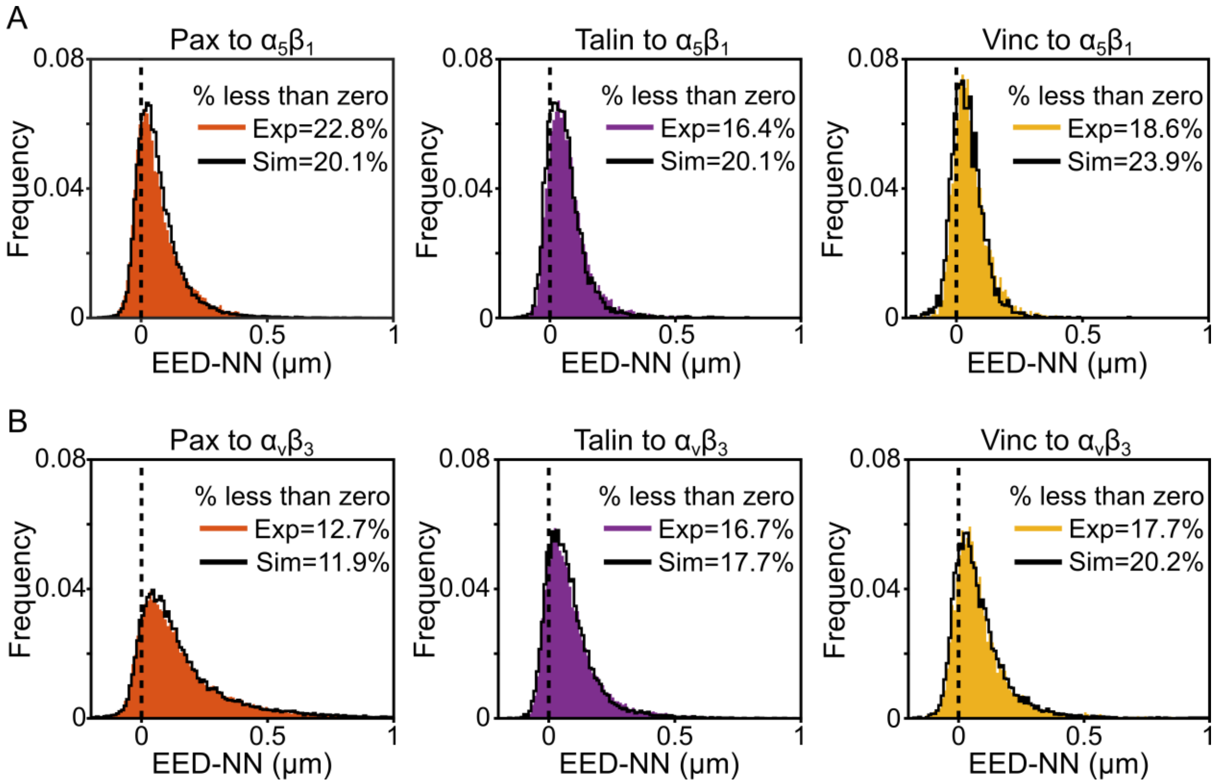


**Figure S6: Edge-to-edge distance for protein nanocluster pairs and percentage of nanoclusters that physically overlap with each other.**

*(A,B) Histogram distribution for EED-NN from adaptors to $\alpha_5\beta_1$ (A) or to $\alpha_v\beta_3$ (B). Solid black lines in each plot (A,B) correspond to histograms for the simulated data sets. Vertical dashed black lines at EED-NN=0 are shown. Values < 0 correspond to overlapping nanoclusters and their percentage of overlapping nanoclusters is indicated in each individual panel. Bin width is 10 nm.*

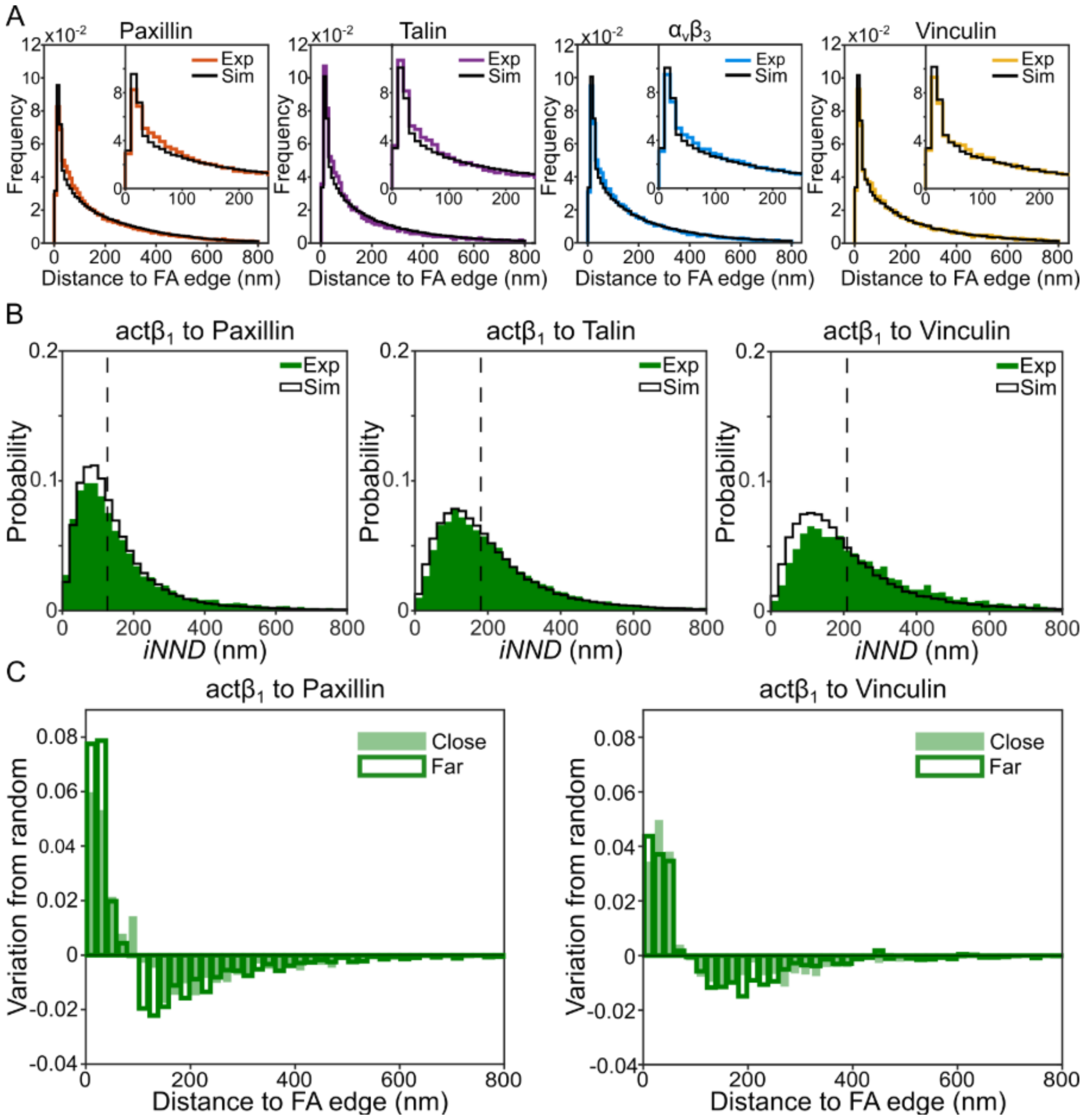


**Figure S7: Lateral organization of nanoclusters inside FAs, with respect to the FA edge.** (A) *Distance to FA edge distributions obtained from DBSCAN-analyzed STORM images. Each histogram represents the frequency of the nanocluster distance-to-edge distribution over all cells. Experimental (Exp) data sets are shown as the colored histograms in each panel, corresponding to paxillin (orange), talin (purple), integrin $\alpha_v\beta_3$ (blue) and vinculin (yellow), whereas the simulated uniform distributions (Sim) are shown as black histograms in each panel. Bin width is 10 nm. (B) Distribution of iNNDs between active $\beta_1$ (act$\beta_1$) integrin nanoclusters and adaptor protein nanoclusters in FAs. Histograms of the iNNDs between nanoclusters of the different proteins pairs as indicated, measured over all cells for both experimental (Exp; shown as solid green bars) and simulated (Sim; shown as black solid lines) random distributions. Dashed vertical lines mark the median distance for each experimental pair. The bin width is 20 nm. (C) Distance to FA edge for act$\beta_1$ nanoclusters close or far from adaptor protein nanoclusters. Variation from random for experimental distance-to-edge plots for act$\beta_1$ nanoclusters in FAs, considering their NND (close vs. far) to paxillin (left) and vinculin (right) nanoclusters. Bin width is 20 nm.*

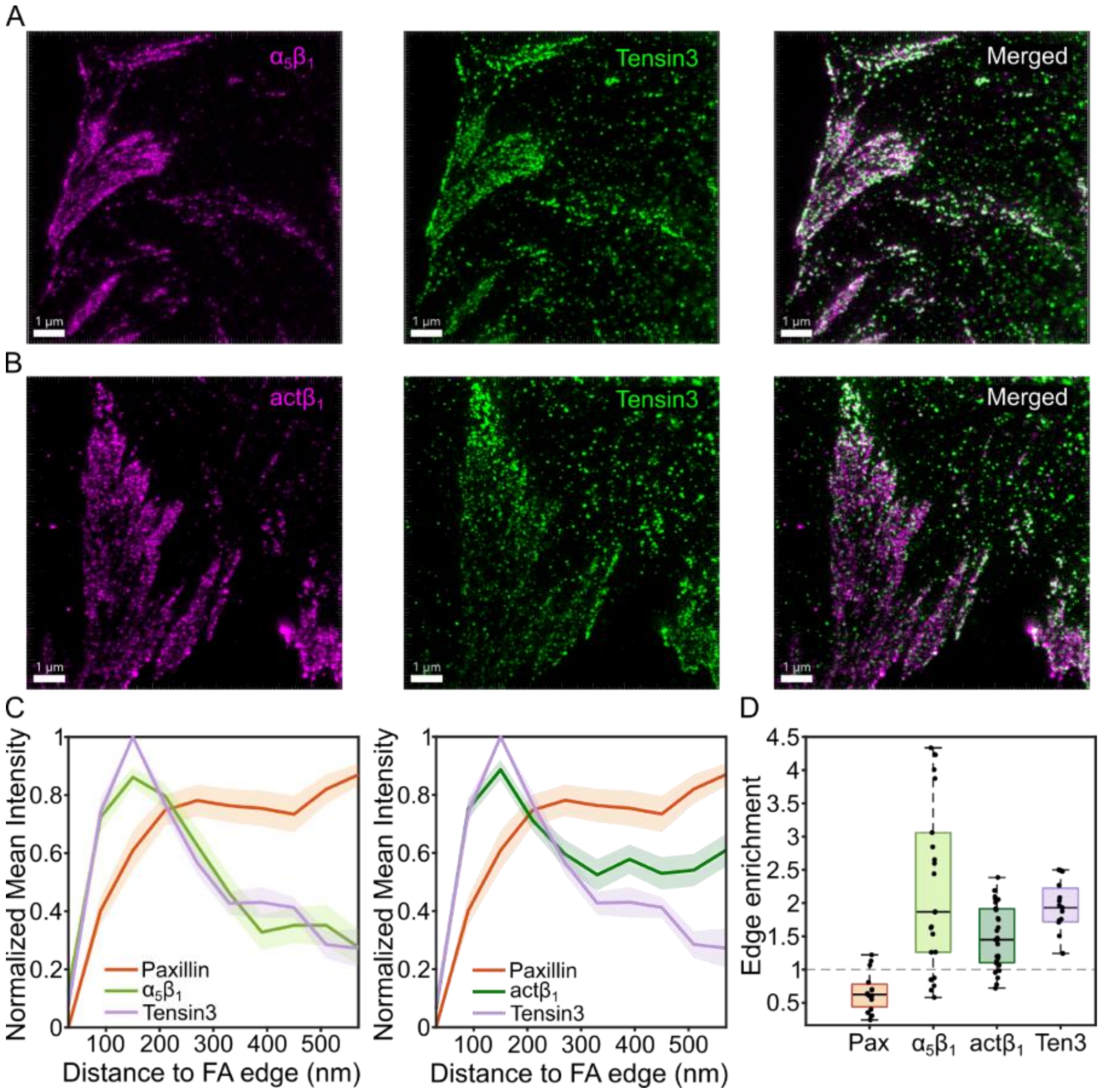


**Figure S8: Dual-color STED reveals radial organization and edge enrichment of β1 integrins and tensin-3 at the FA periphery**. *(A) Dual-color STED images of $\alpha_5\beta_1$ integrin and tensin-3, shown as individual channels and merged views. (B) Dual-color STED images of active $\beta_1$ integrin and tensin-3, shown as individual channels and merged views. (C) Normalized radial mean intensity profiles (mean ± SEM) for $\beta_1$ integrins and tensin-3, computed from the focal adhesion (FA) edge inward up to 600 nm. Individual FA profiles were min-subtracted and normalized to the range [0–1] prior to averaging. (D) Edge enrichment, defined as the ratio of normalized signal in the peripheral FA region (60–180 nm) relative to the FA interior (180–600 nm), is shown for the indicated proteins. The dashed line at 1 denotes equal edge and interior distribution. Paxillin was used as a spatial control. Statistical significance was assessed using two-sided Welch's t-tests comparing Paxillin to $\alpha_5\beta_1$, active $\beta_1$, and tensin-3 (***$p < 0.001$). $N \geq 20$ focal adhesions from multiple cells.*

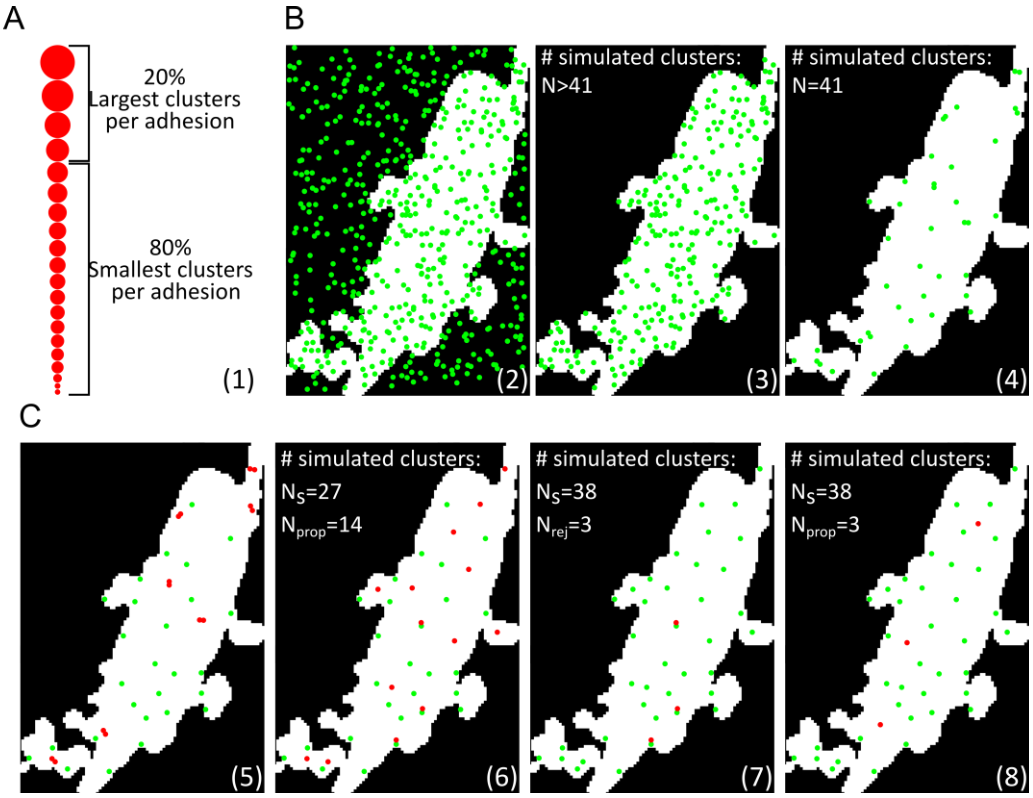


**Figure S9: Generation of random non-overlapping circular nanoclusters in a given adhesion.** *(A) Representation of step 1 of the algorithm where experiment nanoclusters are sorted by size to find the 20% largest nanoclusters, $N_{Big}$. (B) Representation of steps 2–4 of the algorithm where $N_{big}$ points (41 in the example given) are generated and fall inside the FA. (C) Representation of steps 5–8 of the algorithm where it is ensured that none of the $N_{big}$ points overlap with each other. $N_s$ correspond to the points that survive, i.e. do not overlap with any other point, $N_{rej}$ are the points that are rejected because they overlap (red dots) and $N_{prop}$ are newly proposed points. See Supplementary Text 1 for a more detailed explanation.*
